# Experimental Evidence for Phosphorylation-Driven Allosteric Regulation of Alpha Synuclein Function

**DOI:** 10.1101/2025.02.20.639338

**Authors:** Ashlyn N. Dollar, Ian K. Webb

## Abstract

Phosphorylation of serine 129 (pS129) in the intrinsically disordered protein alpha synuclein has long been associated with neurodegenerative disease. In the past several years, the functional relevance of pS219 has been uncovered by electrophysiology, immunoprecipitation, and proteomics as intricately connected with neurotransmitter release and synaptic vesicle (SV) cycling. Unexpectedly, binding to SNARE complex proteins VAMP-2 and synapsin only occurs with phosphorylation-competent alpha synuclein. The VAMP-2 binding domain has been shown to be residues 96-110, which does not include the phosphorylated residue, hinting at allosteric regulation of alpha synuclein protein-protein interactions by pS129. Within this study, cross-linking, covalent labeling, and collision induced unfolding of alpha synuclein and pS129 – as well as an additional encountered form in the brain, oxidized-M1, M5, M116, M127 alpha synuclein – are studied utilizing tandem mass spectrometry. Collision induced unfolding of proteins gives a fingerprint of the structures’ relative compactness and stabilities of various conformations. Covalent labeling of proteins identifies solvent accessible residues and reveals the hydrophobicity (or hydrophilicity) of their microenvironment, while cross-linking of proteins maps the proximity of residue pairs. The combination of collision induced unfolding, covalent labeling, and cross-linking show unequivocally that phosphorylated-S129 alpha synuclein results in a more stable, more compact form. Our results provide evidence of an extensively folded amphipathic region that interacts strongly with the VAMP-2 binding domain. The phosphorylation-induced folding of the amphipathic region likely tunes other protein-protein interactions and interactions with SVs and membranes.

## INTRODUCTION

Alpha synuclein (aSyn) is an intrinsically disordered protein known to form aggregates in patients’ brains correlating with neurological diseases, most notably being Parkinson’s Disease (PD) and Lewy Body (LB) Dementia.^1-3^ aSyn has three distinct regions: the amphipathic/N-terminal region from the N-terminus to residue E61 with helical characteristics, the non-amyloid β component (NAC) region from residues E61 to V95, and the highly acidic region from residue K96 to the C-terminus that is known to be completely disordered.^4,5^

It is known that aSyn undergoes post-translational modifications within the brain, particularly phosphorylation S129 and methionine oxidation on M1, M5, M116, and M127 but there are many discrepancies on how these PTMs affect the structural ensembles of aSyn. 90% of the aggregated alpha synuclein species in those with PD and LBs is found to be phosphorylated S129 (p-aSyn) compared to less than 5% p-aSyn in healthy, post-mortem brain.^2,6,7^ Oxidized M1, M5, M116, and M127 aSyn (ox-aSyn) occurs when the cells within the brain undergo oxidative stress which is linked to PD and LBs.^8,9^ Synucleinopathies may be related to the aSyn species present,^10^ indicating that perhaps oxidized and phosphorylated forms may potentially form more toxic aggregates than unmodified aSyn itself.

While literature has, in the past, suggested that p-aSyn both increases and decreases in toxicity, oligomerization, and change in conformation, the data are conflicting.^11-13^ For example, NMR data have been suggested to be in support of both folding and increases in flexibility of p-aSyn.^14,15^ More recently, electrophysiology measurements of neuron activity show correlations between excitation and the presence of pS129.^16^ Mice knocked in with a S129A mutant, making aSyn phosphorylation incompetent, showed decreased neuronal plasticity.^16^ Low-grade cellular oxidative stress *in vivo* has been shown to promote aSyn aggregation and oligomerization as well, but there’s conflicting literature as to whether oxidized alpha synuclein promotes or inhibits protein fibrillation as well as becomes more stable or unstable than wild type aSyn.^9,17-19^ Another study has also seen that when alpha synuclein is well-folded in the presence of cell crowders like TMAO, the fibrillation is completely inhibited.^20^ Considering these discoveries, the role of phosphorylation on aSyn folding needs to be restudied in light of gain in function conformational changes instead of the formation of aggregates.

Phosphorylation of S129 has been recently revealed to be necessary for proper neuronal function, a carefully localized and controlled regulator of neurotransmission, and an important regulatory binder of the SNARE complex, which controls neuronal signaling in exocytosis via mediation of the fusion of SVs with neuron membranes.^16,21-23^ Specifically, pS129 aSyn has been shown via immunoprecipitation and proteomics to bind VAMP-2, synapsins, and other proteins involved in endo/exocytosis, SV recycling, and vesicle transport pathways, indicating pS129 aSyn as a critical regulator of these processes.^23^ Although pS129 allows these interactions – the S129A mutant disallows them – the interaction sites for VAMP-2 and synapsin are not localized to pS129; deletion variants of aSyn show that the preferred binding site for both VAMP-2 and synapsin is residues 96-110.^23,24^ This led to the proposal of an allosteric mechanism for regulation of these interactions. Though there have been some models of pS129 simulated via artificial intelligence and molecular dynamics approaches,^23,25^ clear experimental evidence of pS129-directed folding of aSyn is needed to validate the allosteric hypothesis.

Similarly, we also studied the fully methionine oxidized form of aSyn, for which the literature also has conflicting data as to whether this species increase or decreases amyloid formation. For example, a coarse grain-MD study demonstrated that ox-aSyn has greater surface area, indicating a higher amount of disorder and exposure to solvent, and that the ox-aSyn oligomers do not form fibrils.^26^ Another study utilizing CD, NMR, and fluorescence anisotropy found that ox-aSyn has a lower affinity for SDS micelles, affecting the physiological function of aSyn, decreasing SV cycling, and increasing the population of aSyn capable of causing toxic oligomers.^27^ IM-MS and CD experiments demonstrated that ox-aSyn has significantly more disorder than aSyn due to a lower propensity of secondary structure, causing it to have less fibrils form due to the oligomer pathways being different between aSyn and ox-aSyn as a result of the methionine oxidation.^28^ Another NMR study that focused on preferential methionine oxidation on aSyn demonstrated that long-range spatial contacts exist between structural segments with M5 and the segments containing either M116 or M127 or even both M116 and M127.^29^

As protein structure defines protein function, it is critical to characterize how the relative PTMs affect aSyn structural ensembles to better understand the function and similarly the pathology of synucleinopathies. Mass spectrometry provides unique specificity in structural measurements with the ability to directly report on the number, sites, and types of PTMs, co-translational modifications, mutations, and truncations (i.e., proteoforms).^30^ This study explores the conformations of aSyn utilizing the various following mass spectrometry techniques: collision induced unfolding,^31^ cross-linking,^32^ and covalent labeling,^33-35^ providing evidence that aSyn is allosterically regulated via S129 phosphorylation.

## METHODS

Please see the Supporting Information (SI) for information about the reagents, alpha synuclein expression and purification, preparation of oxidized alpha synuclein, cross-linking/covalent labeling reaction conditions and concentrations, trypsin digest sample preparation, and oligomerization conditions. All XL/CL experiments were performed in biological buffers and were exchanged into suitable solvents for electrospray or digestion.

### Experimental Design – Mass Spectrometry. Top-Down Approach

Reaction mixtures were diluted to 5μM protein with 250mM ammonium acetate and then electrosprayed into a Waters Synapt G2Si IM-MS (Wilmslow, UK) modified with an ECD cell (Agilent Technologies, Santa Clara, CA). Protein charge states that had a *m/*z shift indicating cross-linking (XL) or covalent labeling (CL) were *m/z* isolated and subjected to electron capture dissociation (ECD) and collision induced dissociation (CID). Low gas pressures – trap gas flow was 10 mL/min, helium cell gas flow was 100 mL/min, IMS gas flow was 20 mL/min - and hot ECD filament settings were used to maximize fragmentation/sequence coverage. 20-minute acquisition times were used for the MS/MS fragmentation experiments. ECD cell settings used for the tandem MS experiments are in SI Table S2a with a constant current set for 2.4A. Trap CID varied depending on the charge state and ranged from 20-55V with constant Transfer CID set to 4V in tandem with the ECD fragmentation for each experiment. Software tools used to view and analyze fragments were ExDViewer v4.6.12 (Agilent Technologies) and Prosight Lite^36^.

The data was collected over three different days with the same reaction conditions (triplicate). We utilize an in-house Python script to verify that fragments are valid and are in at least two out of three replicates. The script then removes potential false positive identifications (i.e., isotopic ion distributions for fragments found in both the cross-linked data and in unmodified datasets). Manual validation with ExDViewer is then performed to validate XL/CL assignments. Our previously published protocol instructions for the Python script utilizing ExDViewer v.4.6.0 remains the same but has been updated to work with ExDViewer v4.6.12. The updated Python script can be found in the SI.

### Bottom-Up Approach

For the bottom-up approach, a Waters ACQUITY UPLC M-Class coupled with a Thermo Scientific Q Exactive Orbitrap Mass Spectrometer was used. The peptide mixtures (XL/CL) were diluted to ~70μg/mL in 250mM ammonium acetate pH 7. 350ng of XL/CL peptides was injected into our Waters ACQUITY UPLC M-Class with a Waters Symmetry® C18, 100Å, 5μm, 300μm x 50mm trap column in tandem with a Waters HSS T3, 100Å, 1.8μm, 300μm x 150 mm C18 column. Solvent A was LC grade water, 0.1% formic acid and solvent B was LC grade acetonitrile, 0.1% formic acid. The sample was loaded onto the trapping column for 2.4min at 10μL/min and then the flow rate was slowed to 4μL/min, both with 95%

A. After loading, the gradient went from 95%-40% A at 4μL/min over 45 minutes and data collection was stopped.

Experiments were conducted in positive mode. For the full MS scan, the automatic gain control (AGC) target was 3e6, maximum IT was 100 ms, and scan range was set to 400-2000 *m/z* with the resolution set to 70,000. The dd-MS^2^ settings was set to have a resolution of 17,500, AGC target 5e5, maximum IT 150ms, TopN 25, isolation window 1.3 *m/z*, and scan range 200-2000 m/z. The stepped normalized collision energy (NCE) was set to 15, 20, and 25 V. The dd settings were set to minimum AGC target of 1e4, intensity threshold 6.7e4, and a dynamic exclusion time of 6 seconds. The data for the bottom-up approach were analyzed using three replicates for each of the XL/CL reaction mixtures. To view and analyze cross-linked and monolinked fragments, pLink/pLabel were used^37^ with a false discovery rate of 2% to determine which peptides were identified to have been modified based on the ions found after the peptide was subjected to HCD fragmentation. Annotations from pLabel were manually verified. To view the CL fragments for both the Gly-Pro experiments and DEPC experiments, pFind/pBuild were used with a false discovery rate of 2% to determine which peptides were identified to have been modified.

### Collision Induced Unfolding Studies

Reaction mixtures were diluted to 5μM protein with 250mM ammonium acetate and then electrosprayed on the Synapt. High gas pressures – trap gas flow was 10 mL/min, helium cell gas flow was 200 mL/min, IMS gas flow was 50 mL/min – and gentle transmission settings were used to maintain compact protein conformations. 2-3 minute acquisition times were used for the ion-mobility (IM) experiments. ExD cell settings used for CIU experiments are given in Table S2b. Trap collision energy was applied prior to injection into the mobility cell to unfold ions. The data was collected over three different days with the same instrument conditions (triplicate). Software tools used to average and visualize the triplicate data sets were MassLynx, TWIMExtract and CIU Suite 2.^38, 39^

## RESULTS AND DISCUSSION

### Intact MS

The electrospray signal was much more stable – generally lasting at least 4 hours for 5μL with static nanoelectrospray– for the ox-aSyn than that of aSyn or p-aSyn. The p-aSyn was much more difficult to stably nanoelectrospray than aSyn, sometimes only lasting 10 minutes before clogging the emitter. This observation is consistent with p-aSyn aggregating more rapidly, at least in ammonium acetate solution, than the other two forms; however, this should not be interpreted as direct evidence that p-aSyn is more aggregation prone in general.

The native charge state distributions (CSD) for the proteoforms of interest are unique to each species (Figure 1). Higher charge states in higher relative abundance are known to correlate with more unfolded structures, and, in general, to disordered and dynamic proteins.^40-46^ The relative abundance of the 9+-15+ charge states of the aSyn and ox-aSyn is much higher than that of the p-aSyn, again indicating that these proteoforms have less structure/less folding than the p-aSyn. p-aSyn has higher relative abundance of the lower charge states (7+ and 8+) than the others; since lower charge states indicate more compactness,^47-49^ it can be hypothesized that the p-aSyn is more compact than the other two proteoforms. Many studies have explored the relationship between IDPs and CSDs, with a consensus stating that IDPs are more sensitive than folded proteins to electrospray conditions and are more prone to variation within replicates.^40,49,50^

**Figure 1.**
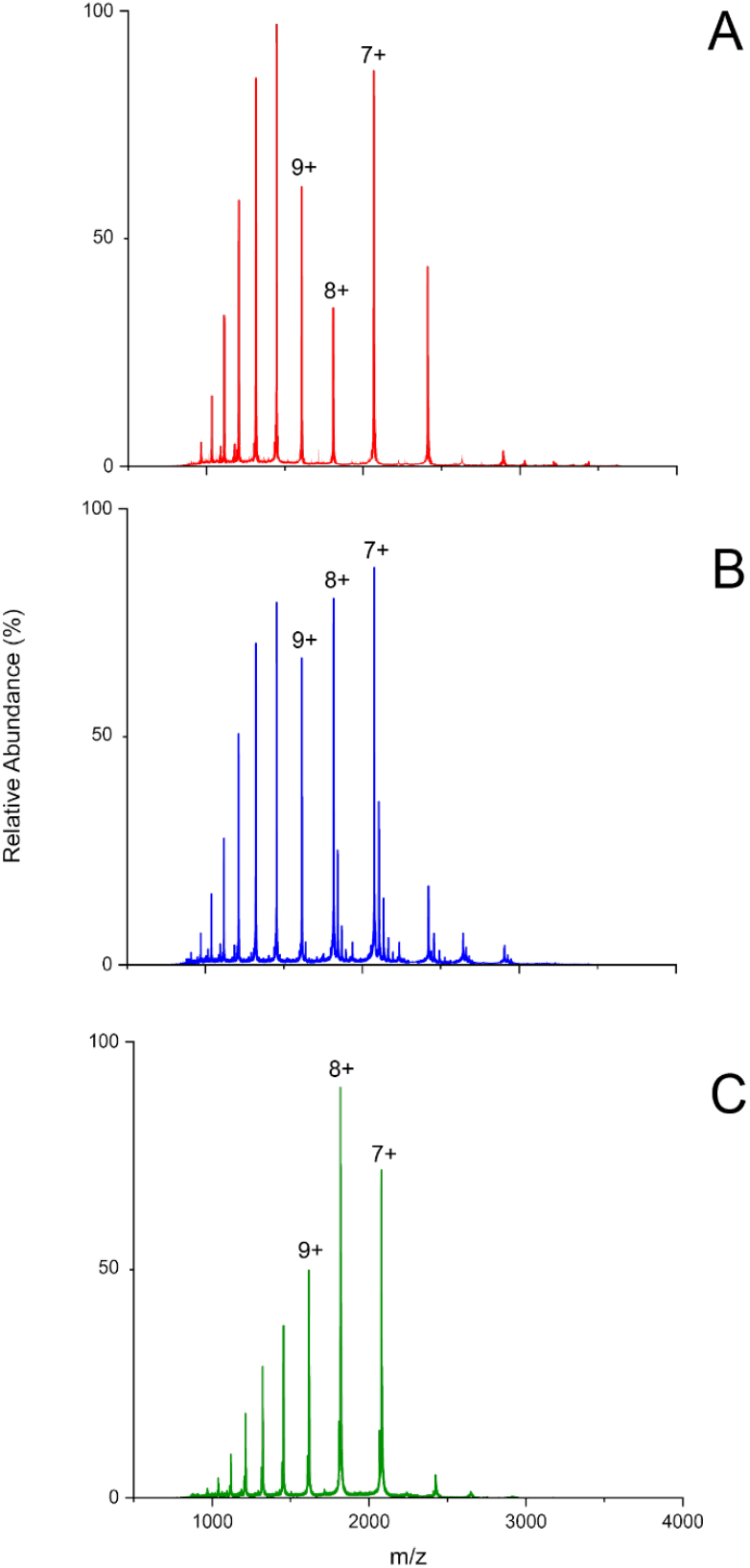
Mass spectra of aSyn (A), oxidized M1, M5, M116, M127 aSyn (B), and phosphorylated S129 aSyn (C) proteoforms under the same native instrument and solution conditions. The 7+ - 9+ charge states are indicated for each.

The CSD implications are in agreement with recent molecular dynamic (MD) studies that find phosphorylation of S129 on aSyn results in a more stable, compact structure for both membrane-bound and in aqueous solution environments, respectively.^25, 51^ The MD studies from Mattaparthi 2024 and De Bryun et. al 2024 do demonstrate that the environment of the phosphorylated proteoform – either membrane-bound or in aqueous solution, respectively – have contradicting results as to whether the phosphorylation of S129 on aSyn inhibits or promotes amyloid formation. The molecular dynamics study of membrane-bound pS129 aSyn showed that pS129 reduces α-strands in the NAC region – the hydrophobic, central region – which is generally known to cause aggregates, indicating that the phosphorylation inhibits amyloid formation in a membrane-bound environment and is more stable than the wild type (WT) aSyn. Modeling of the soluble pS129 aSyn gave evidence that phosphorylation of S129 on aSyn causes the NAC region to form β-hairpin structures, causing aSyn to be more stable and compact, but also more prone to form aggregates in an aqueous environment. Though our study was undertaken in an aqueous environment, our results agree with recent literature that the pS129 results in compaction based on the CSDs above.

### Collision Induced Unfolding

Collision induced unfolding (CIU) is an ion mobility-mass spectrometry (IM-MS) technique that is used to characterize protein stability.^52, 53^ IM is a tool that rapidly (millisecond timescale) separates ions based on their size/shape – their collision cross sections (CCS) – and charge.^54^ More compact ions traverse the IM cell faster than more extended ions. In CIU, the ions undergo energetic collisions, unfolding the protein ions. The ions then enter the IM cell which separates the ions and allows for characterization of their relative stabilities in the gas phase, yielding a so-called CIU “fingerprint”.^31^ CIU is an extremely sensitive technique with a wide range of applications that are, but not limited to, characterizing how PTMs, protein-ligand binding, and mutations in a protein sequence can alter protein structure/structural ensembles.^31^

Previously, tandem CIU experiments have revealed that pS129 aSyn has a higher population of the most compact state in the gas phase than the wild-type.^55^ Figure 2 clearly shows that the unfolding of the 8+ charge states for the three proteoforms have different unfolding patterns, characteristics, and stabilities in the gas phase. Each of the proteoforms have four defining characteristics that have relatively similar drift times in the IM cell. Figure 2 demonstrates that aSyn has the two conformers – indicated by the two different features – merging, indicating that they may be interconverting on the timescale of our experiment (i.e., millisecond or faster transitions).^56^ This is not surprising as aSyn has many structural ensembles being an IDP, so some conformers when unfolded may look like another set of conformers. Strikingly, the p-aSyn only shows significant intensity of the most unfolded structure after the application of collision energy, where the other two proteoforms show all four conformations without adding any energy, indicating that pS129 supports more energetically stable compact structures compared to both aSyn and ox-aSyn.

**Figure 2.**
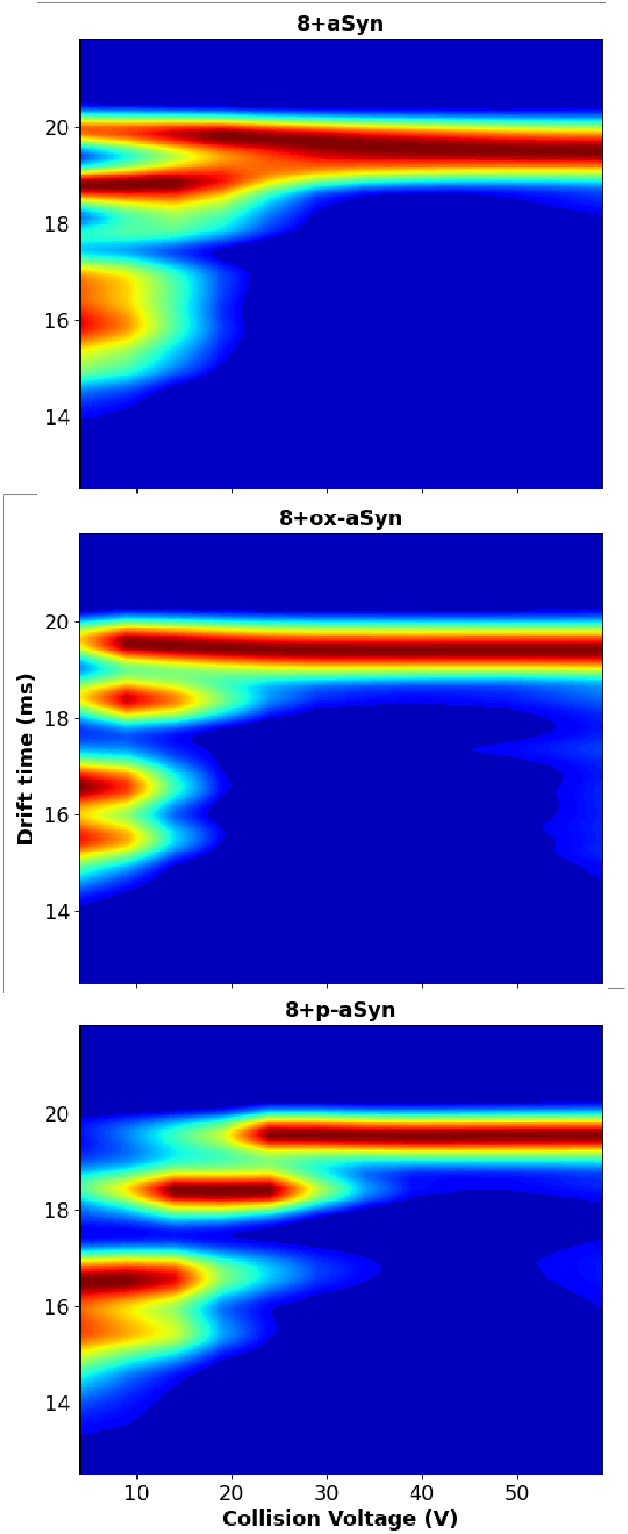
Collision induced unfolding studies on the isolated 8+ charge states for aSyn, oxidized M1, M5, M116, and M127 aSyn, and phosphorylated S129 aSyn where the collision energy applied varied from 0, 5, 10, 15, 20, 25, 30, 40, 50, and 60V within the trap cell. The portrayed CIU plots are averaged using CIU Suite2 for each proteoforms over three replicates completed on different days using the same instrument conditions.

### BS2G and BS3 Cross-linking

Chemical cross-linkers like BS2G and BS3 are traditional, free amine reactive molecular rulers utilized to measure the distance between residues. Cross-linkers are used as a molecular biology tool with many applications, one of which is to characterize protein structure based on intramolecular cross-links identified.^57-59^ Previously, we and others have used cross-linking to characterize the conformational ensemble of aSyn and to characterize changes in conformations upon formation of phase-separated condensates.^60-62^ For this study, BS2G and BS3 were used to identify whether aSyn, ox-aSyn, and p-aSyn have different cross-linking sites due to the presence of PTMs, providing information of how the PTMs affect the overall structure. Both bottom-up and top-down proteomic workflows were employed to determine the cross-linked sites on the aSyn proteoforms of interest as well as characterize the proteoforms.

Bottom-up and top-down proteomics are both mass spectrometry techniques that characterize proteins and these techniques have been compared extensively.^63-65^ The bottom-up workflow proceeds via enzymatic digestion to create peptides. LC-MS/MS then separates and sequences peptides, identifying residues that have modifications like PTMs, XL, and CL. The top-down workflow is when an intact protein is directly introduced – traditionally by electrospray ionization – to the mass spectrometer without enzymatic digestion. MS/MS on the intact protein can then be used to sequence the protein, identify PTMs, XL, and CLs. The lack of intact mass measurement in bottom-up prevents determining whether cross-linked peptides came from a doubly, triply, etc. cross-linked protein. As a result, the first sites to be cross-linked cannot be identified. With the possibility of observing a cross-linked peptide that originates from a protein with multiple cross-links, it is important to remember that cross-links can act as a stabilizer, locking the structure in place, so cross-linking with too much excess linker can potentially disturb the native structure. IM-MS and NMR studies have shown that controlled cross-linking does stabilize a protein slightly while preserving the overall 3-dimensional structure.^60,66^ The top-down workflow is ideal for cross-linking proteoforms studies because the protein with only one or two cross-links and even no cross-links can be isolated and analyzed, so the first, most reactive and/or most favorable cross-link sites can be identified, helping to identify the most critical interactions.

We chose to characterize the three proteoforms with both BS2G and BS3 because they have different spacer arms, allowing us to more thoroughly explore the aSyn conformations than with a single cross-link. BS2G is the shorter spacer arm, 7.7Å, and BS3 has a longer spacer arm, 11.4Å. The cross-linking maps (Figure 3) show that the cross-linked sites are significantly different for each of the aSyn proteoforms. Both the top-down and bottom-up data for aSyn with both cross-linkers demonstrate long-range intermolecular contacts, for example, between lysines 43 and 96, present for both linkers in the top-down and bottom-up data, indicating important interactions between the amphipathic region and the C-terminal region. The ox-aSyn top-down and bottom-up data demonstrate that there is a lack of long-range contacts, and a lack of cross-links at the N-terminus. These results suggest that ox-aSyn is more extended/flexible than a-Syn. The p-aSyn top-down data with BS2G indicates that p-aSyn is more compact in the first 100 residues than either of the other proteoforms, with several identified links between the N-terminus and residue 50, and between 60 and 100. The bottom-up data for the p-aSyn utilizing BS2G, the shorter linker, confirms our findings from the top-down data. The bottom-up data for the p-aSyn linked with BS3, the longer linker, suggests that there are long-range contacts, and that the N-terminal region is well-folded. Figure 3 summarized is that cross-linking the three aSyn proteoforms of interest with two different length cross-linkers is that ox-aSyn is most extended, aSyn shows the presence of some long-range interactions, and p-aSyn is well-folded in the region the cross-links explore (N-terminus to lysine 102). The XL data suggest pS129 stabilizes a more folded form, in agreement with pS129 being the functionally active form and that pS129 is an allosteric regulator of membrane/SV and SNARE complex binding.

**Figure 3.**
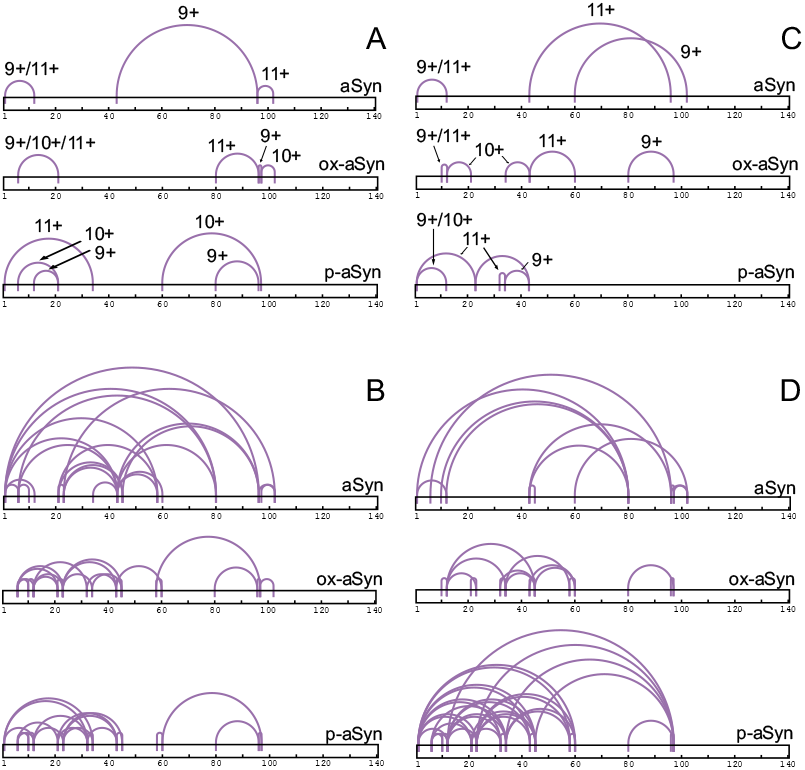
The XLs identified with the top-down approach (A/C) and bottom-up approach (B/D) with aSyn, oxidized M1, M5, M116, M127 aSyn, and phosphorylated S129 aSyn proteoforms using BS2G (A/B) and BS3 (C/D). The charge states are identified for the cross-link identifications determined by the top-down experiments. Sequence ladders for the top-down cross-link identifications for BS2G (A) and BS3 (C) are in SI Figures S3a-b and S4a-b, respectively. A table summarizing the top-down cross-link identifications for BS2G and BS3 is in Figure S5. Tables summarizing bottom-up cross-link identifications utilizing the BS2G (B) and BS3 (D) linkers are in Figures S10 and S11, respectively.

### Gly-Pro Covalent Labeling

Gly-Pro is a heterobifunctional covalent label – labeling both free amines and carboxylic acids on a protein – that utilizes carbodiimide chemistry.^35^ This hydrophilic reagent labeled the three aSyn proteoforms of interest at different residues, demonstrating differences in the solution phase structures of the proteoforms (Figure 4, PDB 1XQ8^67^). The sequence ladders for the 9+, 10+, and 11+ charge states of the three proteoforms within this study labeled with Gly-Pro can be found in the Figure S6a-b and the CLs identified with top-down study are tabulated in Figure S7.

**Figure 4.**
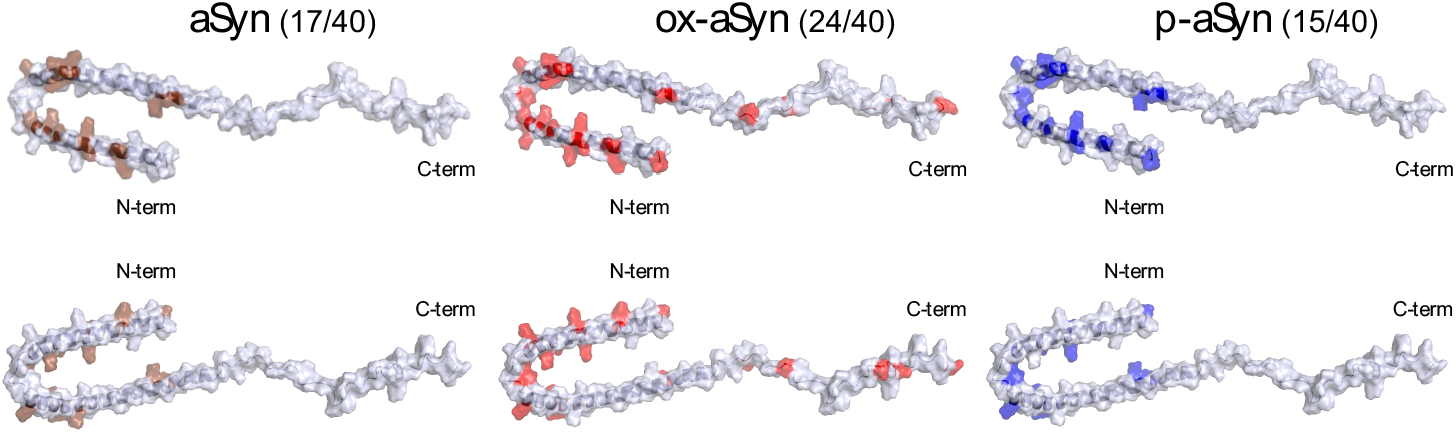
The CLs identified utilizing the bottom-up approach with aSyn, oxidized M1, M5, M116, M127 aSyn, and phosphorylated S129 aSyn proteoforms using cleavable Gly-Pro. The numbers next to the specified proteoforms are how many covalent labels were identified on the protein out of the 40 possible free amine and carboxylic acid residues. A summarized, tabulated version of this data is in SI Figure S12.

The most interesting CLs identified in the top-down experiments are the ones seen at the acidic tail of ox-aSyn. These are CLs in the acidic region of the protein at D119 for the 10+ and E123 for the 11+. As seen with both top-down and bottom-up workflows, aSyn and p-aSyn were not labeled at the C-terminus. As already stated, CLs indicate solvent accessible residues.^68-72^ Therefore, our data potentially suggests that the oxidation is causing the tail to be less compact and/or no longer interact via long-range contacts with the rest of the protein, allowing for this part of the protein to be more solvent accessible. Agreeably, our bottom-up data also demonstrates that the C-terminal tail is Gly-Pro labeled with ox-aSyn and is absent with both aSyn and p-aSyn. Recent in-cell NMR and STED experiments have shown that methionine oxidation on aSyn inhibits fibrillation, so our study agrees that a more exposed/extended tail may play a role in preventing oligomerization.^73^

### DEPC Covalent Labeling

To further characterize the complex relationship that PTMs have on the structural ensembles of aSyn, another CL was used and analyzed with both the top-down and bottom-up approach. DEPC is a widely used CL, known to react strongly with lysine, histidine, free N-termini, and cysteine residues and then weakly with serine, threonine, and tyrosine residues.^70^ Sequence ladders for the top-down experiments for the three aSyn proteoforms reacted with DEPC and the tabulated CL site identifications can be found in Figures S8a-c and S9. Figure 5 shows the CL sites of DEPC identified using the bottom-up proteomics approach for aSyn, ox-aSyn, and p-aSyn. Lysine residues were labeled for all three proteoforms, in agreement with the XL and Gly-Pro labeling that showed that all lysines were accessible and reactive.

**Figure 5.**
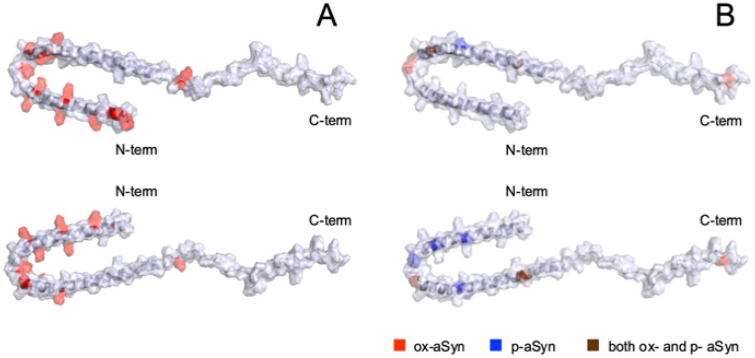
The CLs identified utilizing the bottom-up approach with aSyn, oxidized M1, M5, M116, M127 aSyn, and phosphorylated S129 aSyn proteoforms using DEPC. (A) represents the K/H residues found to have been labeled for all three proteoforms of interest. (B) represents the S/T/Y residues found to have been labeled for all three proteoforms of interest – notably, aSyn did not have any S/T/Y residues labeled. A summarized, tabulated version this bottom-up data is in SI Figure S13.

These data also demonstrated differences in the protein structural ensembles. aSyn was not labeled by DEPC on any S, T, and Y residues. The ox-aSyn data demonstrated only two uniquely labeled S, T, or Y residues, serine 42 (although threonine 44 was labeled in both the oxidized and pS129 forms), and Y136, which provides evidence that the C-terminus is perhaps more accessible or more reactive for the oxidized from (*vide supra*). However, DEPC CLs at T22, T33, Y39, T59, and T64 were all uniquely identified as labeled sites with p-aSyn, demonstrating that the amphipathic region is significantly restructured upon phosphorylation despite the PTM being at S129. DEPC is known to label S, T, and Y residues at more hydrophobic (in other words more folded) microenvironments.^70,74^ Therefore, the DEPC data indicate that there is generally more folding (so perhaps more order) in due to the pS129. It is important to note that the DEPC covalent labeling reaction for p-aSyn required double the concentration of DEPC (8:1 DEPC:protein molar ratio) used with aSyn (4:1 molar ratio) due to significantly slower reaction rates. DEPC generally labels S, T, and Y residues when at higher local concentrations^70^, so there could potentially be a bias for the S, T, and Y labels within the p-aSyn samples due to having a significantly higher final DEPC concentration. However, using a higher concentration for p-aSyn allowed for roughly the same mole ratios of unreacted protein to singly, doubly, triply and quadruply covalently labeled protein as for aSyn (Figure S14). Another reason this potential bias does not seem to be an issue is that while the ox-aSyn had the most S, T, and Y residues labeled, this reaction mixture contained the lowest concentration of DEPC (2:1 molar ratio) required to reach the similar degrees of labeling to the other two forms (Figure S14).

Due to the three proteoforms requiring different DEPC concentrations, we also explored the kinetics for the DEPC reactions which have been shown to be primarily a function of higher order structure, with slower rates attributed to more folded structures due to the lack of surface accessibility of reactive side chains.^34^ Dose response curves (Figure S15) were used to calculate second-order rate constants for DEPC labeling. The rate constant for reaction with p-aSyn was over one order of magnitude lower than the rate constants for ox-aSyn and aSyn (3.4 ± 0.1 × 10^−5^, 1.4 ± 0.1 × 10^−4^, and 1.36 ± 0.08 × 10^−4^ μM^−1^s^−1^, respectively). DEPC labeling kinetics provide additional support that phosphorylation at S129 causes significant compaction, likely stabilizing a binding-competent and thus functional state.

## CONCLUSIONS

The biophysical mass spectrometry techniques employed in this study– CIU, cross-linking, and covalent labeling – provides experimental evidence for significant folding of pS129 aSyn and supports the hypothesis that pS129 is an allosteric regulator of aSyn function, tuning its structure for binding SNARE proteins and membranes/SVs. The CIU experiments showed that unfolding p-aSyn requires more collision energy in the gas phase while ox-aSyn and aSyn required similar but less energy for unfolding. Cross-linking of the three proteoforms with two different length cross-linkers demonstrated p-aSyn has a more well-folded amphipathic region. Cross-linking also demonstrated that aSyn does have some long-range interactions while ox-aSyn does not, demonstrating that the ox-aSyn has the most extended structures. Covalent labeling with both hydrophilic and hydrophobic cross-linkers again show that ox-aSyn has more extended conformers due to the acidic tail of the protein being labeled. Covalent labeling of p-aSyn showed extensive labeling of tyrosine and threonine in the amphipathic region, providing additional evidence of folding. Our work provides structural evidence that pS129 activates aSyn for its functional roles via conformational stabilization and a structural mechanism for the biological function of pS129 determined via previous electrophysiological and immunoprecipitation studies.

## Supporting information

Python Script for Analysis of Top Down Data from Viewer

## SUPPORTING INFORMATION

Additional information about reagents, protein expression and purification, cross-linking and covalent labeling reaction conditions, trypsin digestion conditions, and data for both top-down and bottom-up experiments (PDF)

Updated script for data verification (Python script)

## ACKNOWLEDGEMENTS

This work was funded by the National Institute of General Medical Sciences of the National Institutes of Health under R35GM151251 (IKW).

