## Supporting Information for "Experimental Evidence for Phosphorylation-Driven Allosteric Regulation of Alpha Synuclein Function"

### Exploring the Conformationome of Alpha Synuclein and its Proteoforms with Various Mass Spectrometry Techniques

**Table of Contents**

*Additional information about:*


Reagents

Protein expression and purification

Preparation of oxidized alpha synuclein

Crosslinking/covalent labeling reactions

Trypsin digest

*Figures:*

S1. Oxidized aSyn Sequence Ladders

S2. ExD Cell Settings

a. ECD combined with CID experiments
b. CIU experiments

S3. Bis(sulfosuccinimidyl) glutarate (BS2G) Crosslinking Sequence Ladders

a. oxidized M1, M5, M116, M127 aSyn (ox-aSyn)
b. phosphorylated S129 aSyn (p-aSyn)

S4. Bis(sulfosuccinimidyl) suberate (BS3) Crosslinking Sequence Ladders

a. ox-aSyn
b. p-aSyn

S5. Table of BS2G and BS3 Crosslinking Sites Identified with Top-Down MS/MS for 9+-11+ Charge States

S6. Glycyl-*L*-proline (Gly-Pro) Covalent Labeling Sequence Ladders

a. ox-aSyn
b. p-aSyn

S7. Table of Gly-Pro Covalent Labeling Sites Identified with Top-Down MS/MS for 9+-11+ Charge States

S8. Diethyl pyrocarbonate (DEPC) Covalent Labeling Sequence Ladders

a. aSyn
b. ox-aSyn
c. p-aSyn

S9. Table of DEPC Covalent Labeling Sites Identified Utilizing Top-Down MS/MS for 9+-11+ Charge States

S10. Table of BS2G Crosslinking Sites Identified Utilizing Bottom-Up (Trypsin Digestion) LCMS/MS

S11. Table of BS3 Crosslinking Sites Identified Utilizing Bottom-Up (Trypsin Digestion) LCMS/MS

S12. Table of Gly-Pro Covalent Labeling Sites Identified Utilizing Bottom-Up (Trypsin Digestion) LCMS/MS

S13. Table of DEPC Covalent Labeling Sites Identified Utilizing Bottom-Up (Trypsin Digestion) LCMS/MS

S14. Mass Spectra of DEPC Covalent Labeling with Equal Ratios of Labeled Product

S15. DEPC Covalent Labeling Kinetics

**Reagents.**

Ammonium acetate, acetonitrile, and tris-base were purchased from Fisher Scientific (Waltham, MA). 3(N-morpholino)propane sulfonic acid (MOPS), bis(sulfosuccinimidyl) glutarate (BS2G), and bis(sulfosuccinimidyl) suberate (BS3) was purchased from ThermoFisher Scientific (Waltham, MA). Glycyl-L-proline (Gly-Pro) was obtained from Tokyo Chemical Industry (Tokyo, Japan). 1-ethyl-3-(3-dimethylaminopropyl)carbodiimide (EDC) was purchased from Santa Cruz Biotechnology (Dallas, TX). 1-hydroxy-7-azabenzotriazole (HOAt) was obtained from Apex Biotechnology (Glen Allen, VA). Diethyl pyrocarbonate (DEPC), N-methylpyrrolidine (NMP), and phosphate buffered saline (PBS) were purchased from Sigma Aldrich (St. Louis, MO). Hydrogen peroxide, 35% wt was obtained from Acros Organics, now ThermoFisher Scientific (Waltham, MA). Bio-spin columns with bio-gel P-6 in tris-X buffer with a 6kDa cutoff were used from buffer exchange from BioRad (Hercules, CA). S129-phosphorylated alpha synuclein (p-aSyn) was purchased from Proteos (Kalamazoo, MI). Trypsin Gold Standard was purchased from Promega (Madison, WI).

**Protein Expression and Purification.**

pET21a-alpha-synuclein was a gift from Michael J Fox Foundation MJFF (Addgene plasmid # 51486 ; http://n2t.net/addgene:51486 ; RRID:Addgene_51486). The untagged aSyn was purified using a previously published protocol.[1] Using a Micro BCA assay, the purified recombinant aSyn was found to be 210µM in 50mM ammonium acetate, pH 7.

**Preparation of Oxidized Alpha Synuclein.**

The reaction conditions to oxidize each methionine (M1, M5, M116, and M127) on aSyn were 200µM aSyn, 320mM hydrogen peroxide in 200mM tris-base pH 7.4 for 15 minutes at room temperature. The reaction was quenched by buffer-exchange two times into 100mM MOPS pH 6.5. Verification of oxidation at each methionine was confirmed doing top-down mass spectrometry analysis for the 7+-12+ charge states which can be found in the SI Figure S1.

**Crosslinking/Covalent Labeling Reactions.**

*BS2G and BS3 Crosslinking Reactions.*

The BS2G or BS3 XLing final reaction concentrations were 50µM protein (of p-aSyn or oxidized aSyn), 150µM crosslinker (BS2G or BS3) in 1X PBS. The reaction went for 15 minutes at room temperature and was quenched with buffer-exchange two times into 250mM ammonium acetate.

*Glycyl-L-proline Covalent Labeling Reaction.*

The glycyl-L-proline CLing final reaction concentrations were 50µM protein (of p-aSyn or oxidized aSyn), 18mM EDC, 20mM HOAT dissolved in NMP, 150mM glycyl-L-proline in 100mM MOPS pH 6.5. The reaction went for 12 minutes at room temperature and was quenched with buffer-exchange two times into 250mM ammonium acetate.

*DEPC Covalent Labeling Reaction for Top-Down and Bottom-Up Analysis.*

The diethyl pyrocarbonate (DEPC) CLing reactions were for 1 minute at 37℃ in 100mM MOPS pH 6.85. These reactions have been studied greatly and were altered slightly from the previously published protocol.[2] It is important to note that the DEPC stock solution was made fresh for each experiment at 20mM in ACN. The reaction was quenched with buffer-exchange into 250mM ammonium acetate. The final reaction concentration was 50μM protein, with final concentrations of DEPC for aSyn to be 200μM, for ox-aSyn to be 100μM, and p-aSyn to be 400μM.

*DEPC Covalent Labeling Kinetics Reactions.*

The DEPC CLing reactions for were for 1 minute at 37℃ in 100mM MOPS pH 6.85. The final reactions were 10μM protein with varying concentrations of DEPC at either 2, 5, 10, 20, 30, 50, 70, 100, 200, or 300 μM. The reactions were quenched by dilution to 5μM protein in 250mM ammonium acetate and then buffer-exchanged into 250mM ammonium acetate to wash away remaining DEPC and MOPS buffer.

**Trypsin Digest.**

For confirmation of XL and CL sites for the multiple aSyn proteoforms, tryptic digests were used. To do this, the crosslinking reactions were completed first as described above, then buffer-exchanged into 50mM tris-X pH 7.4. Trypsin was then added to a ratio of roughly 1:40 protease: protein and allowed to incubate at 37℃ overnight. Formic acid was added to be 1% of the reaction mixture to quench digestion.

S1. Oxidized aSyn Sequence Ladders

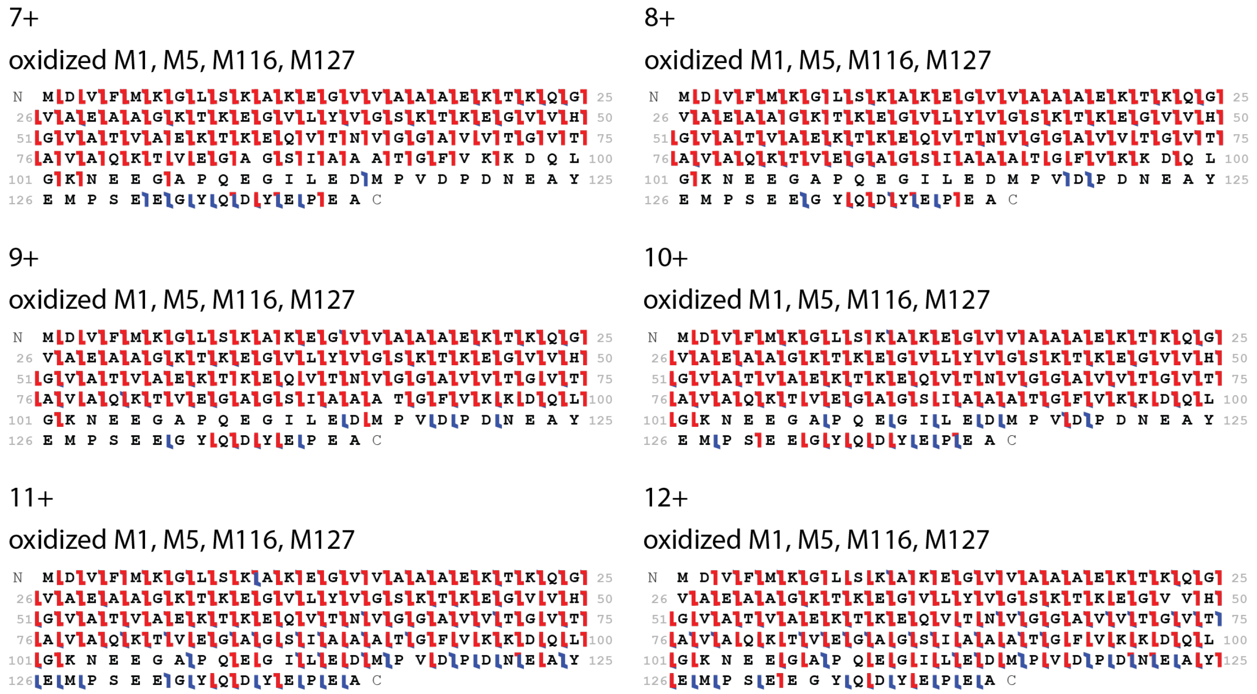


**Figure S1**. Sequence ladders for the 7+-12+ triplicates for oxidized M1, M5, M116, M127 aSyn verifying that the aSyn oxidation reaction does modify each methionine.

#### S2. ExD Cell Settings

a. ECD combined with CID experiments

| L1 | L2 | LM3 | L4 | FB | LM5 | L6 | L7 |
|---|---|---|---|---|---|---|---|
| 0.0 | 0.0 | 6.4 | 7.5 | 4.0 | 7.5 | 10 | 15.0 |

b. CIU experiments

| L1 | L2 | LM3 | L4 | FB | LM5 | L6 | L7 |
|---|---|---|---|---|---|---|---|
| -16.0 | -13.0 | -5.6 | 6.4 | 2.9 | 7.0 | 8.0 | 9.0 |

**Figure S2.** ExD cell settings – displayed as voltages – utilized for the indicated experiments. “L” means lens; “LM” means lens magnet; “FB” means filament bias.

#### S3. Bis(sulfosuccinimidyl) glutarate (BS2G) Crosslinking Sequence Ladders

a. oxidized M1, M5, M116, M127 aSyn (ox-aSyn)

9+

unmodified

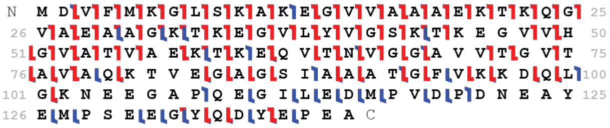


modified

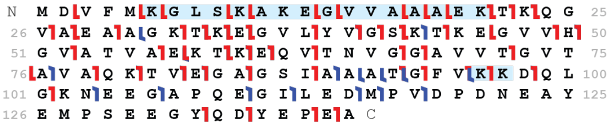


10+

unmodified

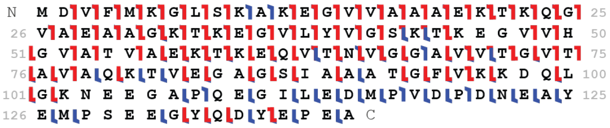


modified

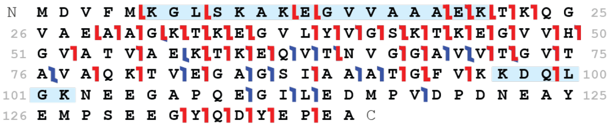


11+

unmodified

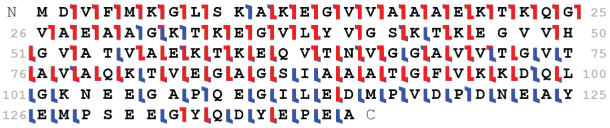


modified

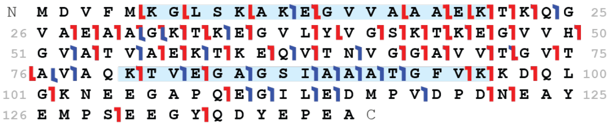


b. phosphorylated S129 aSyn (p-aSyn)

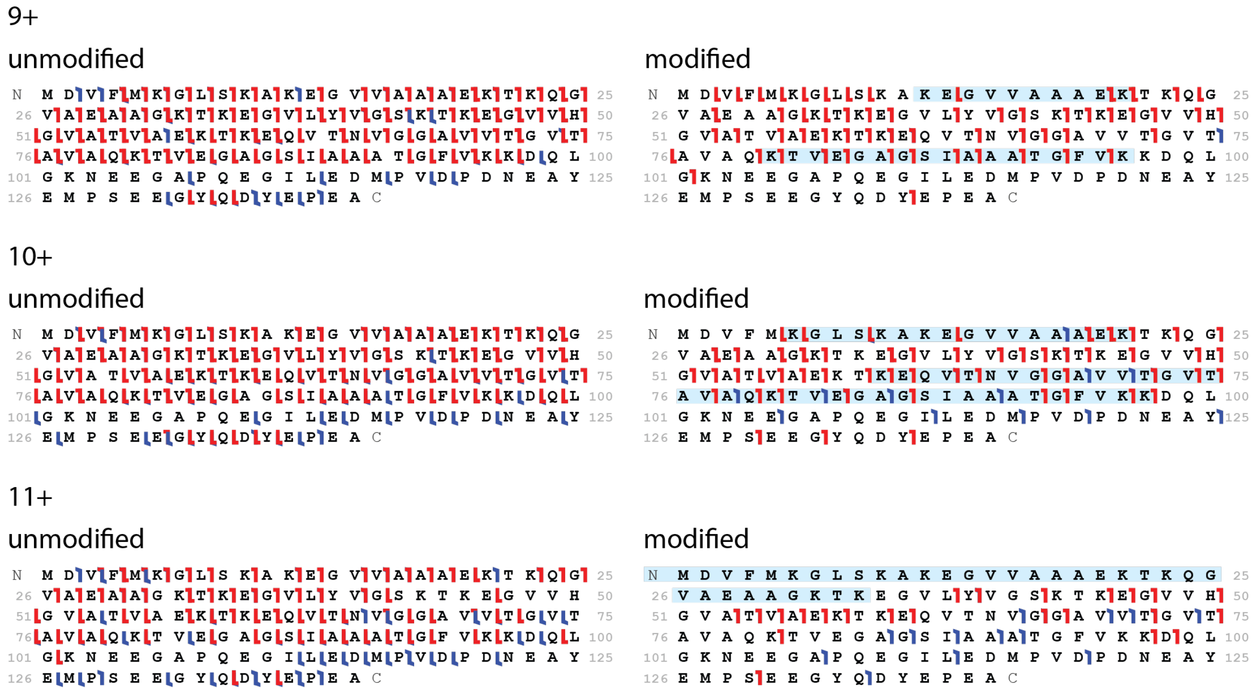


**Figure S3**. Sequence ladders for the 9+-11+ triplicates for ox-aSyn (a) and p-aSyn (b) crosslinked with BS2G.

#### S4. Bis(sulfosuccinimidyl) suberate (BS3) Crosslinking Sequence Ladders

##### a. oxidized M1, M5, M116, M127 aSyn (ox-aSyn)

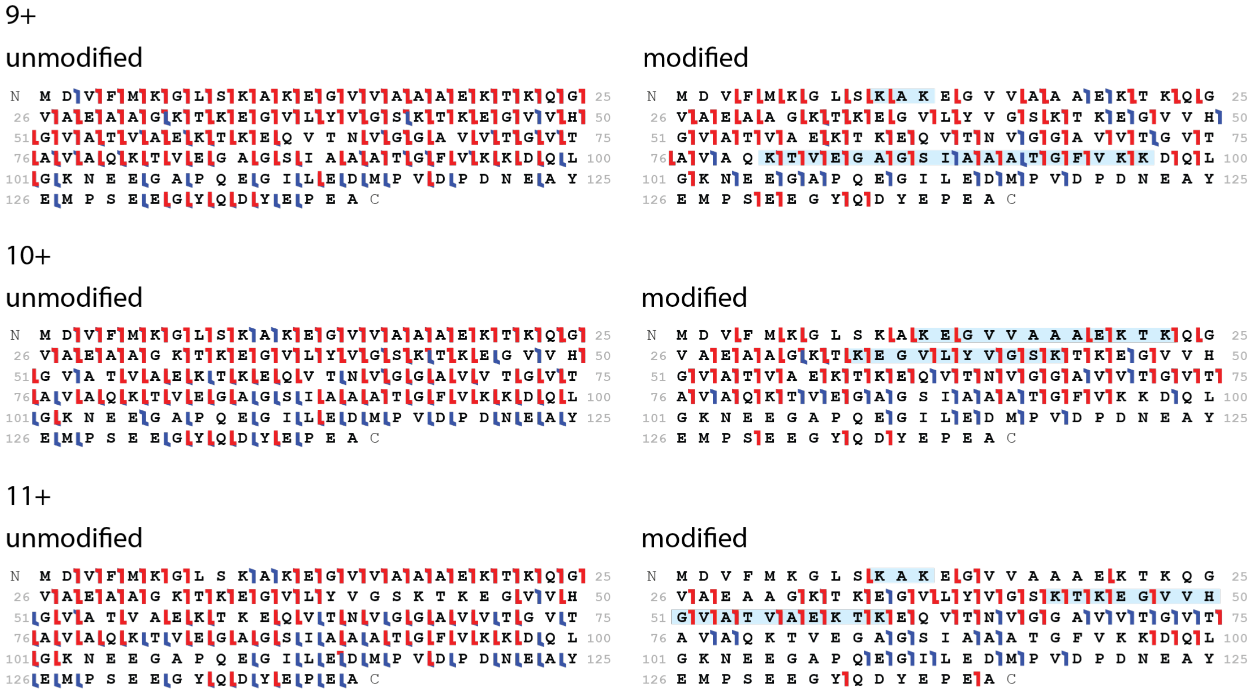


##### b. phosphorylated S129 aSyn (p-aSyn)

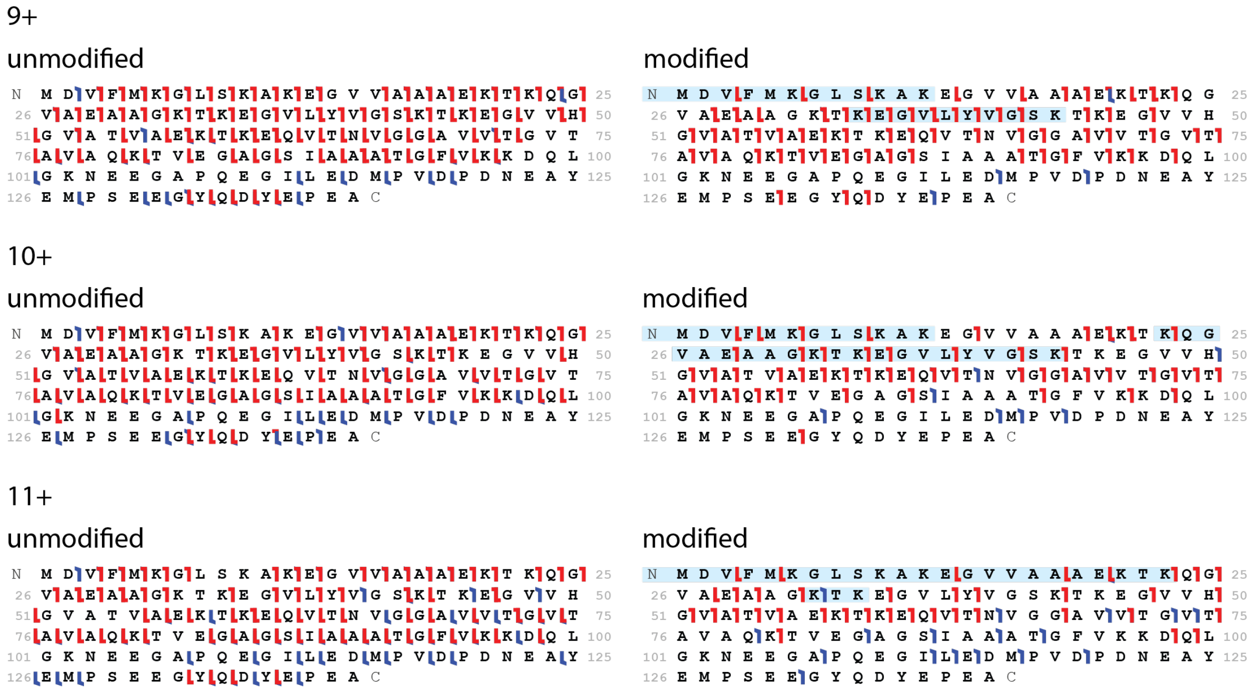


**Figure S4.**. Sequence ladders for the 9+-11+ triplicates for ox-aSyn (a) and p-aSyn (b) crosslinked with BS3.

S5. Table of BS2G and BS3 Crosslinking Sites Identified with Top-Down MS/MS for 9+-11+ Charge States

| | BS2G | | | BS3 | | |
|---|---|---|---|---|---|---|
| | 9+ | 10+ | 11+ | 9+ | 10+ | 11+ |
| aSyn[4] | Nt-K12<br>K43-K96 | - | Nt-K12<br>K96-K102 | Nt-K12<br>K60-K102 | - | Nt-K12<br>K43-K96 |
| ox-aSyn | K6-K21<br>K96-K97 | K6-K21<br>K97-K102 | K6-K21<br>K80-K96 | K10-K12<br>K80-K97 | K12-K21<br>K34-K43 | K10-K12<br>K43-K60 |
| p-aSyn | K12-K21<br>K80-K96 | K6-K21<br>K60-K97 | Nt-K34 | Nt-K12<br>K34-K43 | Nt-K12<br>K23-K43 | Nt-K23<br>K32-K34 |

**Figure S5.** Crosslink sites identified using the native top-down approach for the three proteoforms of interest utilizing the BS2G and BS3 crosslinkers.

S6. Glycyl-*L*-proline (Gly-Pro) Covalent Labeling Sequence Ladders

a. oxidized M1, M5, M116, M127 aSyn (ox-aSyn)

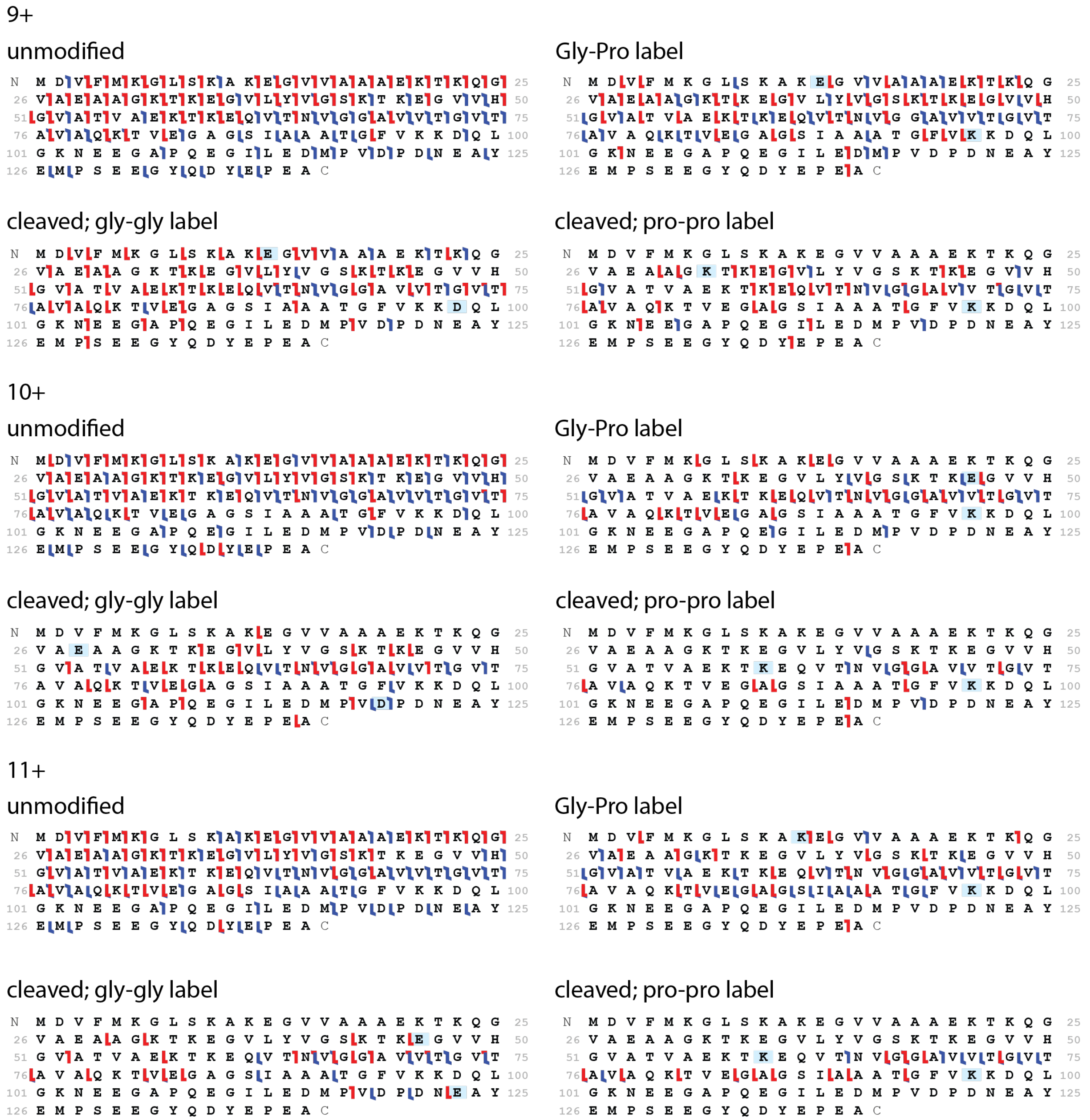

b. phosphorylated S129 aSyn (p-aSyn)

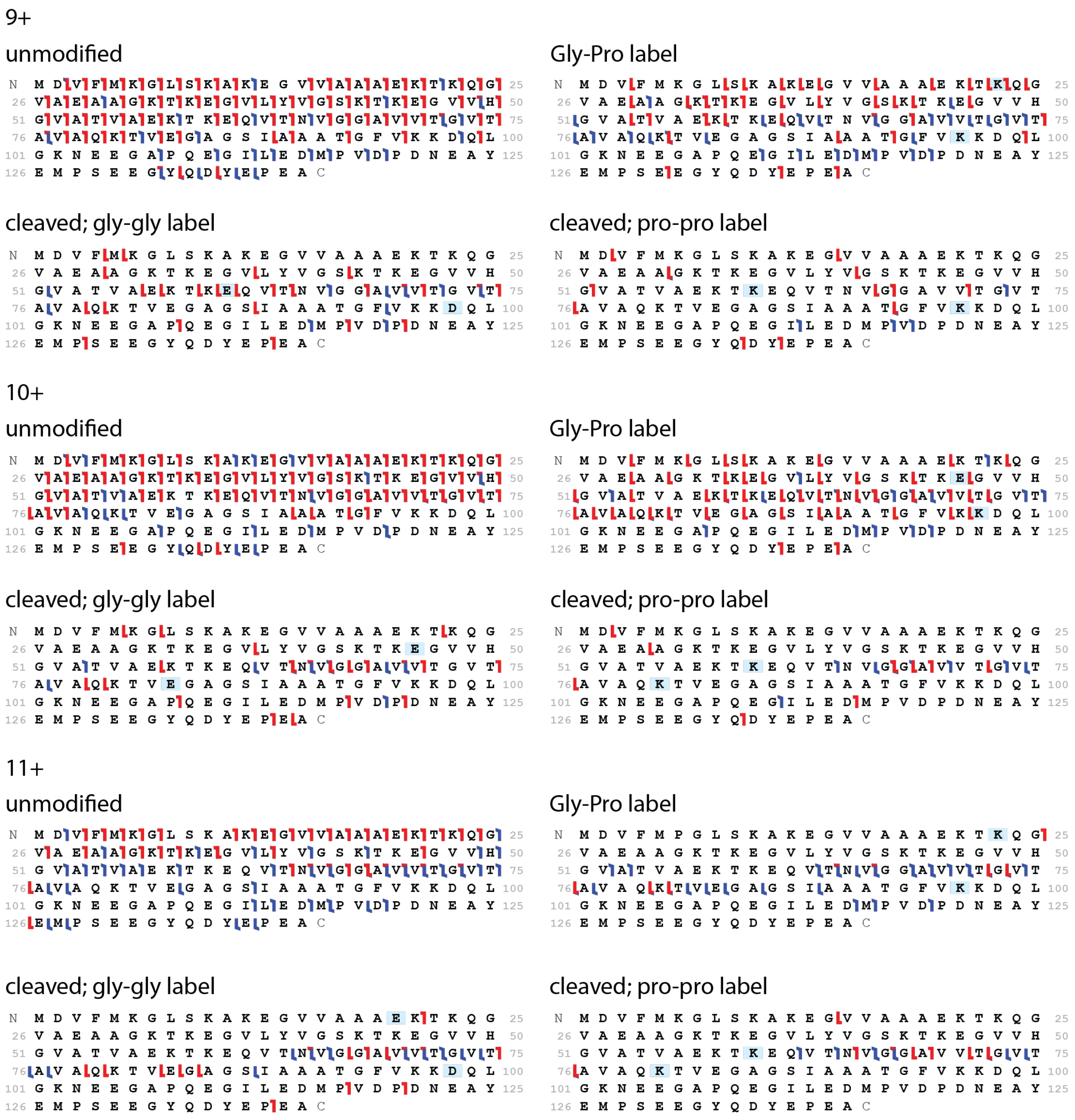


**Figure S6**. Sequence ladders for the 9+-11+ triplicates for ox-aSyn (a) and p-aSyn (b) CLd with cleavable Gly-Pro.

S7. Table of Gly-Pro Covalent Labeling Sites Identified with Top-Down MS/MS for 9+-11+ Charge States

| | | **Covalent label unique to proteoform** | | | **Covalent label found in ≥2 proteoforms** | | | |
|---|---|---|---|---|---|---|---|---|
| | | **aSyn** | **ox-aSyn** | **p-aSyn** | **aSyn and ox-aSyn** | **aSyn and p-aSyn** | **ox-aSyn and p-aSyn** | **aSyn, ox-aSyn, and p-aSyn** |
| **9+** | Glycyl-L-proline CL, Uncleaved | E61, E83 | E13 | K23 | - | - | K96 | - |
| | Glycine, cleaved addition | E83 | E13 | - | - | E61 | D98 | - |
| | Proline, cleaved addition | K80 | K32 | - | - | K60 | K96 | - |
| **10+** | Glycyl-L-proline CL, Uncleaved | E13 | - | - | - | - | E46 | K96/K97 |
| | Glycine, cleaved addition | D2, D98 | E23, D119 | E46 | - | E83 | - | - |
| | Proline, cleaved addition | K12 | K96 | - | - | K80 | K60 | - |
| **11+** | Glycyl-L-proline CL, Uncleaved | - | - | K23 | K12 | - | - | K96 |
| | Glycine, cleaved addition | E61 | E123 | E20 | E46 | D98 | - | - |
| | Proline, cleaved addition | K23 | K96 | K80 | - | - | - | K60 |

**Figure S7**. Table summarizing the CL sites of cleavable Gly-Pro identified with the top-down method for the 9+-11+ triplicates on aSyn, oxidized M1, M5, M116, M127 aSyn, and phosphorylated S129 aSyn.

S8. Diethyl pyrocarbonate (DEPC) Covalent Labeling Sequence Ladders

a. aSyn

9+

unmodified

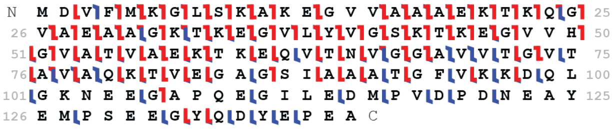


modified

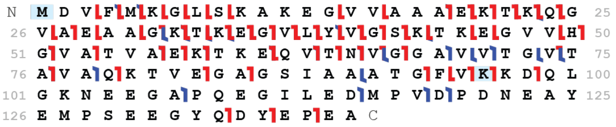


10+

unmodified

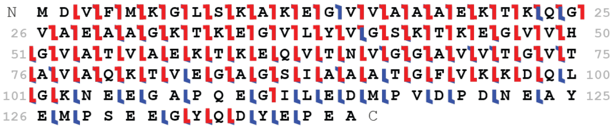


modified

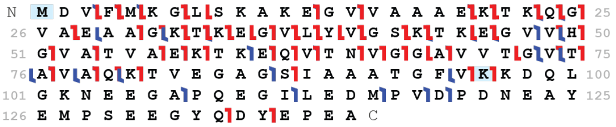


11+

unmodified

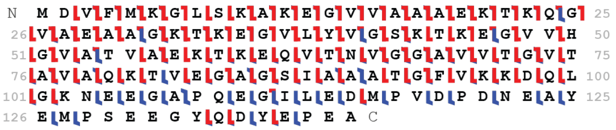


modified

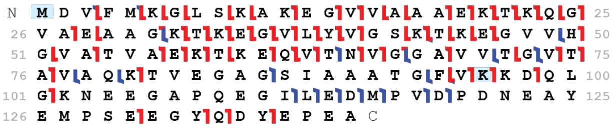


b. oxidized M1, M5, M116, M127 aSyn (ox-aSyn)

9+

unmodified

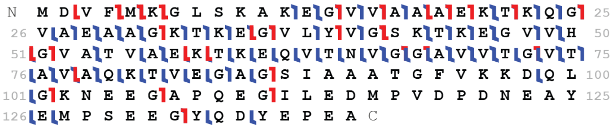


modified

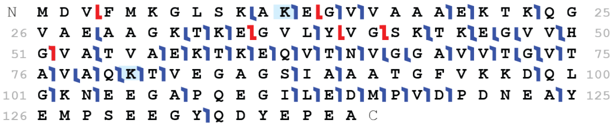


10+

unmodified

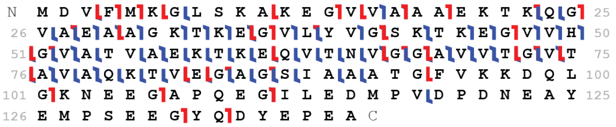


modified

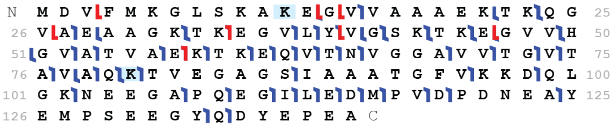


11+

unmodified

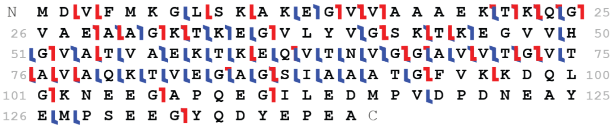


modified

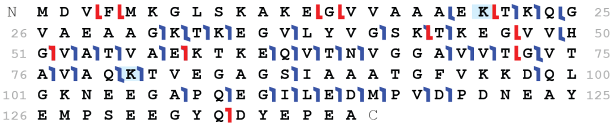

c. phosphorylated S129 aSyn (p-aSyn)

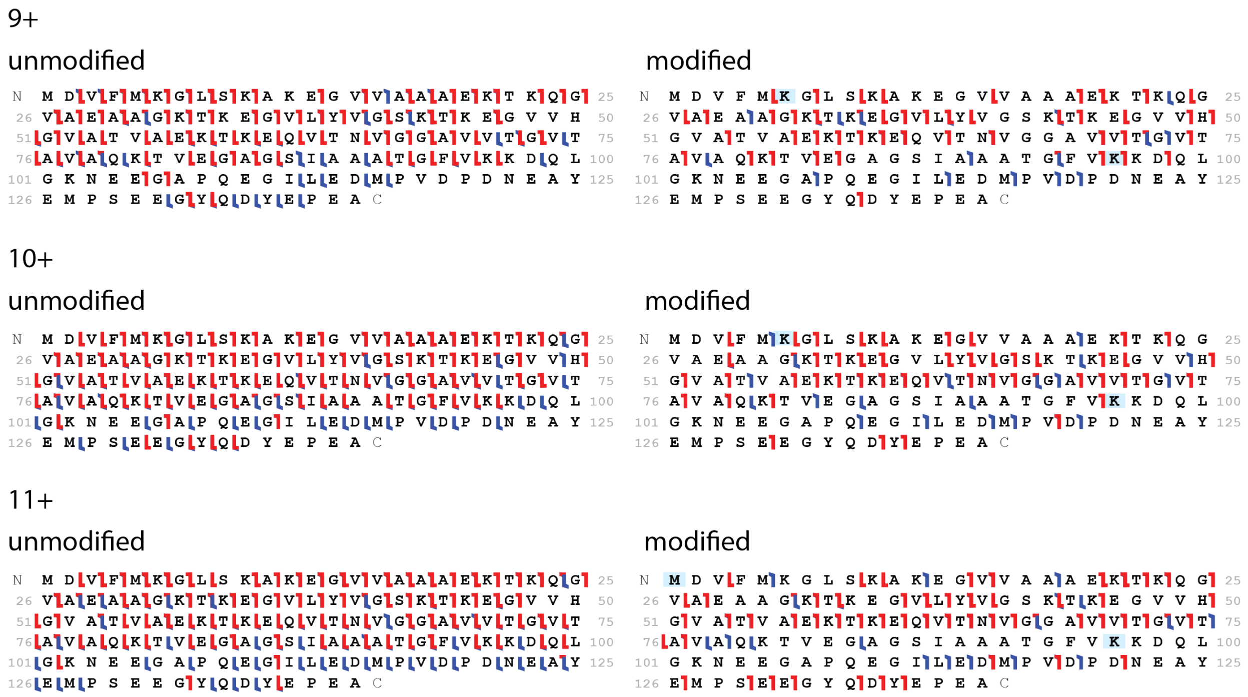


**Figure S8**. Sequence ladders for the 9+-11+ triplicates for aSyn (a), ox-aSyn (b), and p-aSyn (c) CLd with DEPC.

S9. Table of DEPC Covalent Labeling Sites Utilizing Top-Down MS/MS on the 9+-11+ Charge States

| | **9+** | **10+** | **11+** |
|---|---|---|---|
| **aSyn** | Nt, K96 | Nt, K96 | Nt, K96 |
| **ox-aSyn** | K12, K80 | K12, K80 | K21, K80 |
| **p-aSyn** | K6, K96 | K6, K96 | Nt, K96 |

**Figure S9.** The CLs identified utilizing the top-down approach with aSyn, oxidized M1, M5, M116, M127 aSyn, and phosphorylated S129 aSyn proteoforms using DEPC for the 9+, 10+, and 11+ charge states.

S10. Table of BS2G Crosslinking Sites Utilizing Bottom-Up (Trypsin Digestion) LCMS/MS

| Crosslinks unique to proteoform | | | Crosslinks found in ≥2 proteoforms | | | |
|---|---|---|---|---|---|---|
| aSyn | ox-aSyn | p-aSyn | aSyn and ox-aSyn | aSyn and p-aSyn | ox-aSyn and p-aSyn | aSyn, ox-aSyn, and p-aSyn |
| Nt-K6<br>Nt-K43<br>Nt-K58<br>Nt-K80<br>Nt-K96<br><br>K6-K43<br>K6-K80<br><br>K21-K45<br>K21-K58<br><br>K23-K102<br><br>K34-K43<br><br>K43-K58<br>K43-K80<br>K43-K96<br><br>K45-K96<br><br>K96-K102 | K6-K23<br><br>K12-K21<br><br>K34-K45<br><br>K58-K97 | Nt-K32<br>Nt-K34<br><br>K21-K32<br>K21-K43<br><br>K32-K34<br><br>K60-K97 | K43-K60<br><br>K97-K102 | Nt-K12 | K6-K12<br>K6-K21<br><br>K10-K12<br><br>K12-K23<br>K12-K32<br><br>K23-K34<br><br>K32-K43<br><br>K43-K45<br><br>K58-K60<br><br>K80-K96<br><br>K96-K97 | K6-K10<br><br>K21-K23<br><br>K23-K43<br>K23-K45 |

**Figure S10**. Table summarizing the crosslinked residues using BS2G identified with the bottom-up method on aSyn, oxidized M1, M5, M116, M127 aSyn, and phosphorylated S129 aSyn.

S11. Table of BS3 Crosslinking Sites Utilizing Bottom-Up (Trypsin Digestion) LCMS/MS

| Crosslinks unique to proteoform | | | Crosslinks found in ≥2 proteoforms | | | |
|---|---|---|---|---|---|---|
| aSyn | ox-aSyn | p-aSyn | aSyn and ox-aSyn | aSyn and p-aSyn | ox-aSyn and p-aSyn | aSyn, ox-aSyn, and p-aSyn |
| Nt-K80<br>K6-K96<br>K10-K80<br>K12-K80<br>K43-K80<br>K43-K96<br>K60-K102<br>K96-K102<br>K97-K102 | K32-K45<br>K34-K45<br>K43-K60<br>K45-K60 | Nt-K21<br>Nt-K23<br>Nt-K32<br>Nt-K34<br>Nt-K43<br>Nt-K45<br>Nt-K58<br>Nt-K60<br>K6-K10<br>K6-K12<br>K6-K21<br>K6-K34<br>K12-K32<br>K12-K43<br>K12-K97<br>K21-K32<br>K21-K45<br>K23-K43<br>K23-K45<br>K23-K58<br>K23-K60<br>K23-K97<br>K34-K60<br>K34-K97<br>K45-K97 | None | Nt-K12 | K10-K12<br>K12-K23<br>K12-K34<br>K12-K45<br>K21-K23<br>K32-K34<br>K34-K58<br>K58-K60<br>K80-K97 | K43-K45<br>K96-K97 |

**Figure S11**. Table summarizing the crosslinked residues using BS3 identified with the bottom-up method on aSyn, oxidized M1, M5, M116, M127 aSyn, and phosphorylated S129 aSyn.

#### S12. Table of Gly-Pro Covalent Labeling Sites Identified Utilizing Bottom-Up (Trypsin Digestion) LCMS/MS

| Glycyl-L-proline CL, uncleaved | | | Glycine addition, cleaved | | | Proline addition, cleaved | | |
|---|---|---|---|---|---|---|---|---|
| aSyn | ox-aSyn | p-aSyn | aSyn | ox-aSyn | p-aSyn | aSyn | ox-aSyn | p-aSyn |
| | Nt | Nt | E20 | E20 | E20 | | Nt | |
| D2 | D2 | D2 | | E28 | | | K21 | K21 |
| K12 | K12 | | | E57 | | K23 | | K23 |
| E13 | E13 | E13 | | E61 | | | K32 | |
| E20 | E20 | E20 | | D98 | | K34 | K34 | |
| K21 | K21 | K21 | | | | | K45 | |
| | K23 | | | | | | K58 | |
| E28 | E28 | E28 | | | | | K60 | |
| K32 | K32 | | | | | | K80 | |
| | K34 | | | | | | | |
| E35 | E35 | E35 | | | | | | |
| | K43 | K43 | | | | | | |
| E46 | E46 | E46 | | | | | | |
| E57 | E57 | E57 | | | | | | |
| K58 | K58 | K58 | | | | | | |
| K60 | | | | | | | | |
| E61 | E61 | E61 | | | | | | |
| K80 | | K80 | | | | | | |
| E83 | E83 | E83 | | | | | | |
| | D98 | | | | | | | |
| | E104 | | | | | | | |
| | D121 | | | | | | | |
| | E126 | | | | | | | |
| | E139 | | | | | | | |

**Figure S12**. Table summarizing the covalent labeled residues using cleavable Gly-Pro identified with the bottom-up method on aSyn, oxidized M1, M5, M116, M127 aSyn, and phosphorylated S129 aSyn.

#### S13. Table of DEPC Covalent Labeling Sites Identified Utilizing Bottom-Up (Trypsin Digestion) LCMS/MS

| aSyn | ox-aSyn | p-aSyn |
|---|---|---|
| Nt | Nt | Nt |
| K6 | K6 | K6 |
| | K10 | K10 |
| K12 | K12 | K12 |
| K21 | K21 | K21 |
| | | T22 |
| K23 | K23 | K23 |
| K32 | K32 | K32 |
| | | T33 |
| K34 | K34 | K34 |
| | | Y39 |
| | S42 | |
| K43 | K43 | K43 |
| | T44 | T44 |
| K45 | K45 | K45 |
| H50 | H50 | H50 |
| | T54 | T54 |
| K58 | K58 | K58 |
| | | T59 |
| K60 | K60 | K60 |
| | | T64 |
| K80 | K80 | |
| | T81 | T81 |
| K96 | K96 | K96 |
| | K102 | |
| | Y136 | |

**Figure S13**. Table summarizing the covalent labeled residues using DEPC identified with the bottom-up method on aSyn, oxidized M1, M5, M116, M127 aSyn, and phosphorylated S129 aSyn.

#### S14. Mass Spectra of DEPC Covalent Labeling with Equal Ratios of Labeled Product

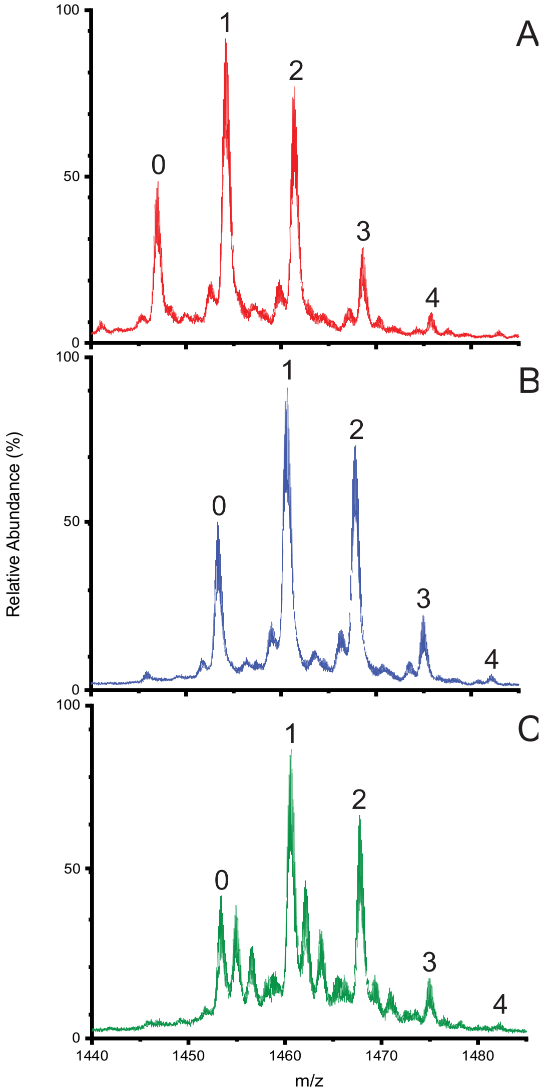


**Figure S14.** Mass spectra of the complex DEPC CL reaction mixture of the 10+ charge states for aSyn (A), ox-aSyn (B), and p-aSyn (C) with the reaction conditions as described in the *DEPC Covalent Labeling Reaction for Top-Down and Bottom-Up Analysis* section of the SI. The numbers above peaks indicate number of CLs.

S15. DEPC Covalent Labeling Kinetics.

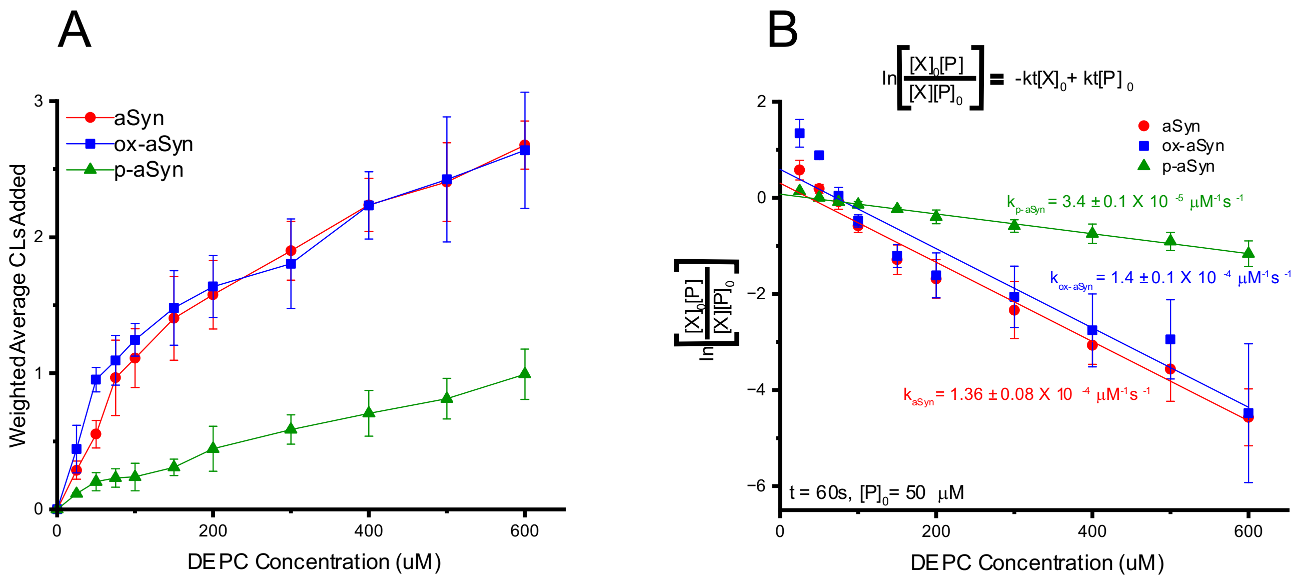


**Figure S15.** (A) Weighted average addition of DEPC covalent label as a function of DEPC concentration. (B) Dose-response curve for the depletion of the unreacted aSyn proteoform by reaction with DEPC, where P refers to unreacted aSyn and X refers to DEPC. In B, [P]/[$P_0$] was estimated by dividing the peak area for the unreacted aSyn by the sum of the peak areas for reacted and unreacted aSyn. [X] = [$P_0$] - [P]. This treatment assumes second-order reaction rates.
