## Supplementary material for "Experimental Evidence for Phosphorylation-Driven Allosteric Regulation of Alpha Synuclein Function": Python Script for Analysis of Top Down Data from Viewer

from sys import exit
import os
import pandas as pd
import csv


cwd = os.getcwd()
group = os.listdir(cwd)

for files in group:
    if files.endswith("old.csv"):
        os.remove(files)
    elif files.endswith("log.txt"):
        os.remove(files)

residue_masses = {
'A': 71.037114,
    'R': 156.101111,
    'N': 114.042927,
    'D': 115.026943,
    'C': 103.009185,
    'E': 129.042593,
    'Q': 128.058578,
    'G': 57.021464,
    'H': 137.058912,
    'I': 113.084064,
    'L': 113.084064,
    'K': 128.094963,
    'M': 131.040485,
    'F': 147.068414,
    'P': 97.052764,
    'S': 87.032028,
    'T': 101.047679,
    'W': 186.079313,
    'Y': 163.06332,
    'V': 99.068414
}

proton_mass = 1.00728
NH4_mass = 18.03383
H3O_mass = 19.01784
H2O_minus_NH_mass = 2.999116

def merge_duplicate_keys(dictionary):
    merged_dict = {}
    for key, value in dictionary.items():
        if key in merged_dict:
            merged_dict[key] += value
        else:
            merged_dict[key] = value
    return merged_dict

def write_deconv_qpi_file(IDs,filename):
    with open(filename+'.csv', 'w', newline='') as file:
        rep_writer = csv.writer(file)
        rep_writer.writerow(["Monoisotopic_Mass", "Sum_Intensity", "Number_of_Charge_States", "Average_Charge", "Deconvoluted_Mass", "Relative_Abundance", "Fractional_Abundance", "RT_Range", "Apex_RT"])
        for line in IDs:
            newline = [line[0],line[1],"1","1","1","1","1","1","1"]
            rep_writer.writerow(newline)

def sum_intensity_isotopes(data_csv_file):
    # Define the input CSV file name
    input_file = data_csv_file

    # Rename the input CSV file by appending '_old'
    os.rename(input_file, input_file.replace('.csv', '_old.csv'))

    # Read the CSV file into a DataFrame
    df = pd.read_csv(input_file.replace('.csv', '_old.csv'))
    
    # For compatability with viewer beta 4.6.12 Need to change monoisotopic weight to "Monoisotopic", and set it to yes if it is the smallest of the theo m/z having the same other ids
    df = df.rename(columns={'Monoisotopic Weight':'Monoisotopic'})
    
    # Convert all numerical inputs to numbers, remove commas from intensity values.
    df['Charge'] = pd.to_numeric(df['Charge'])
    df['Pos'] = pd.to_numeric(df['Pos'])
    df['Theo MZ'] = pd.to_numeric(df['Theo MZ'])
    
    # For compatability with viewer beta 4.6.12
    df['Monoisotopic'] = pd.to_numeric(df['Monoisotopic'])
    
    if df['Intensity'].dtype == 'object':
        df['Intensity'] = df['Intensity'].str.replace(',', '').astype(float)

    # Group by Charge, Type, and Pos and sum Intensity for rows where Monoisotopic is 'Yes'
    df['Intensity'] = df.groupby(['Charge', 'Type', 'Pos', 'Monoisotopic'])['Intensity'].transform('sum')
    
    #F ilter rows for only monoisotope
    df = df.loc[df.groupby(['Charge', 'Type', 'Pos', 'Monoisotopic', 'Intensity'])['Theo MZ'].idxmin()]
    
    # For compatability with viewer beta 4.6.12
    df['Monoisotopic'] = 'Yes'

    # Write the updated DataFrame back to the input CSV file
    df.to_csv(input_file, index=False)
        

def process_list(input_list):
    # Dictionary to store sums associated with unique first items
    sums_dict = {}

    # Iterate through the input list
    result_list = []
    current_first_item = None
    current_sum = 0
    for item in input_list:
        first_item, second_item = item[0], item[1]
        
        # If first item changes, add a new row with the sum
        if first_item != current_first_item:
            if current_first_item is not None:
                result_list.append([current_first_item, current_sum])
            current_first_item = first_item
            current_sum = 0
        
        # Update the sum
        current_sum += second_item
        
        # Store the sum associated with the first item
        sums_dict[first_item] = current_sum

    # Add the last sum
    if current_first_item is not None:
        result_list.append([current_first_item, current_sum])

    return result_list
            
def import_csv_files(csv_files):
    dataframes = []
    for file_path in csv_files:
        try:
            df = pd.read_csv(file_path)
            dataframes.append(df.values.tolist())
            print(f"File {file_path} successfully imported.")
        except Exception as e:
            print(f"Error importing file {file_path}: {e}")
    return dataframes

if __name__ == "__main__":
    script_dir = os.path.dirname(os.path.realpath(__file__))
    csv_files = []
    file_names = ['mod_rep1.csv', 'unmod_rep1.csv', 'mod_rep2.csv', 'unmod_rep2.csv', 'mod_rep3.csv', 'unmod_rep3.csv']
    
    for file_name in file_names:
        file_path = os.path.join(script_dir, file_name)
        if os.path.isfile(file_path):
            csv_files.append(file_path)
        else:
            print(f"File {file_name} not found in the directory.")

    if len(csv_files) < 6:
    
        print("Please make sure all required files are present in the directory.")
        exit(1)

 # Importing the first line of a specific named text file as a string
    with open('protein.txt', 'r') as file:
        first_line = file.readline().strip()
        next_line = file.readline().strip()
        parts1 = next_line.split(':')
        second_line = parts1[0]
        num_charge_neut = int(parts1[1])
        modded_residues = parts1[2].split(';')
        if 'Nt' in modded_residues:
            num_charge_neut_nterm = int(num_charge_neut) -1
        else:
            num_charge_neut_nterm = int(num_charge_neut)
        other_lines = file.readlines()
        static_mods = {}
        for line in other_lines:
            line = line.strip()
            parts = line.split(':')
            number = parts[0]
            values = parts[1].split(';')
            for value in values:
                static_mods[value] = number
                 

    # Creating an int variable set to the length of the string
    string_length = len(first_line)

    data_arrays = import_csv_files(csv_files)


    
mzb_unmod = proton_mass
mzc_unmod = NH4_mass
mzy_unmod = H3O_mass
mzz_unmod = H2O_minus_NH_mass

mzb_mod = proton_mass + float(second_line)
mzc_mod = NH4_mass + float(second_line)
mzy_mod = H3O_mass + float(second_line)
mzz_mod = H2O_minus_NH_mass + float(second_line)

    # Define the filename
filename = 'potential_unmods.csv'
filename1 = 'potential_mods.csv'
filename2 = 'no_statics_unmods.csv'
filename3 = 'no_statics_mods.csv'

# Create lists to hold the rows of the CSV
rows = []
mod_rows = []
no_statics_rows = []
no_statics_mod_rows = []


# Generate rows based on specified ranges
for i in range(1, string_length):
    residue = first_line[i-1]
    b_res_num = str(i)
    b_number = static_mods.get(b_res_num) 
    frag_seq_b = first_line[:i]
    max_charge_b = frag_seq_b.count('K') + frag_seq_b.count('H') + frag_seq_b.count('R') +1
    
            
    if b_number is not None:
        b_numb_flot = float(b_number)
        if residue in residue_masses:
            mzb_unmod = b_numb_flot+ mzb_unmod + residue_masses[residue]
    else:
        if residue in residue_masses:
            mzb_unmod = mzb_unmod + residue_masses[residue]
    rows.append([f'b{i}',mzb_unmod,max_charge_b])
    
    
for j in range(1, string_length):
    residue = first_line[j-1]
    c_res_num = str(j)
    c_number = static_mods.get(c_res_num)
    frag_seq_c = first_line[:j]
    max_charge_c = frag_seq_c.count('K') + frag_seq_c.count('H') + frag_seq_c.count('R') +1
    
    if c_number is not None:
        c_numb_flot = float(c_number)
        if residue in residue_masses:
            mzc_unmod = c_numb_flot + mzc_unmod + residue_masses[residue]
    else:
        if residue in residue_masses:
            mzc_unmod = mzc_unmod + residue_masses[residue]
    next_residue = first_line[j]
    if next_residue == "P":
        rows.append([f'c{j}','',max_charge_c])
    else:
        rows.append([f'c{j}',mzc_unmod,max_charge_c])

for k in range(1, string_length):
    residue = first_line[string_length-k]
    y_res_num = str(string_length+1-k)
    y_number = static_mods.get(y_res_num)
    backward_seq = ''.join(reversed(first_line))
    frag_seq_y = backward_seq[:k]
    max_charge_y = frag_seq_y.count('K') + frag_seq_y.count('H') + frag_seq_y.count('R') +1
    
    if y_number is not None:
        y_numb_flot = float(y_number)
        if residue in residue_masses:
            mzy_unmod = mzy_unmod + y_numb_flot + residue_masses[residue]
    else:
        if residue in residue_masses:
            mzy_unmod = mzy_unmod + residue_masses[residue]
    rows.append([f'y{k}',mzy_unmod,max_charge_y])
    
for l in range(1, string_length):
    residue = first_line[string_length-l]
    z_res_num = str(string_length+1-l)
    z_number = static_mods.get(z_res_num)
    backward_seq = ''.join(reversed(first_line))
    frag_seq_z = backward_seq[:l]
    max_charge_z = frag_seq_z.count('K') + frag_seq_z.count('H') + frag_seq_z.count('R') +1
    
    if z_number is not None:
        z_numb_flot = float(z_number)
        if residue in residue_masses:
            mzz_unmod = mzz_unmod + z_numb_flot + residue_masses[residue]
    else:
        if residue in residue_masses:
            mzz_unmod = mzz_unmod + residue_masses[residue]
    if residue == 'P':
        rows.append([f'z{l}','',max_charge_z])
    else:
        rows.append([f'z{l}',mzz_unmod,max_charge_z]) 
        
# Generate no statics rows based on specified ranges
mzb_unmod = proton_mass
mzc_unmod = NH4_mass
mzy_unmod = H3O_mass
mzz_unmod = H2O_minus_NH_mass
for i in range(1, string_length):
    residue = first_line[i-1]
    b_res_num = str(i)
    b_number = 0 
    frag_seq_b = first_line[:i]
    max_charge_b = frag_seq_b.count('K') + frag_seq_b.count('H') + frag_seq_b.count('R') +1
    
            
    if b_number is not None:
        b_numb_flot = float(b_number)
        if residue in residue_masses:
            mzb_unmod = b_numb_flot+ mzb_unmod + residue_masses[residue]
    else:
        if residue in residue_masses:
            mzb_unmod = mzb_unmod + residue_masses[residue]
    no_statics_rows.append([f'b{i}',mzb_unmod,max_charge_b])
    
    
for j in range(1, string_length):
    residue = first_line[j-1]
    c_res_num = str(j)
    c_number = 0
    frag_seq_c = first_line[:j]
    max_charge_c = frag_seq_c.count('K') + frag_seq_c.count('H') + frag_seq_c.count('R') +1
    
    if c_number is not None:
        c_numb_flot = float(c_number)
        if residue in residue_masses:
            mzc_unmod = c_numb_flot + mzc_unmod + residue_masses[residue]
    else:
        if residue in residue_masses:
            mzc_unmod = mzc_unmod + residue_masses[residue]
    next_residue = first_line[j]
    if next_residue == "P":
        no_statics_rows.append([f'c{j}','',max_charge_c])
    else:
        no_statics_rows.append([f'c{j}',mzc_unmod,max_charge_c])

for k in range(1, string_length):
    residue = first_line[string_length-k]
    y_res_num = str(string_length+1-k)
    y_number = 0
    backward_seq = ''.join(reversed(first_line))
    frag_seq_y = backward_seq[:k]
    max_charge_y = frag_seq_y.count('K') + frag_seq_y.count('H') + frag_seq_y.count('R') + 1
    
    if y_number is not None:
        y_numb_flot = float(y_number)
        if residue in residue_masses:
            mzy_unmod = mzy_unmod + y_numb_flot + residue_masses[residue]
    else:
        if residue in residue_masses:
            mzy_unmod = mzy_unmod + residue_masses[residue]
    no_statics_rows.append([f'y{k}',mzy_unmod,max_charge_y])
    
for l in range(1, string_length):
    residue = first_line[string_length-l]
    z_res_num = str(string_length+1-l)
    z_number = 0
    backward_seq = ''.join(reversed(first_line))
    frag_seq_z = backward_seq[:l]
    max_charge_z = frag_seq_z.count('K') + frag_seq_z.count('H') + frag_seq_z.count('R') +1
    
    if z_number is not None:
        z_numb_flot = float(z_number)
        if residue in residue_masses:
            mzz_unmod = mzz_unmod + z_numb_flot + residue_masses[residue]
    else:
        if residue in residue_masses:
            mzz_unmod = mzz_unmod + residue_masses[residue]
    if residue == 'P':
        no_statics_rows.append([f'z{l}','',max_charge_z])
    else:
        no_statics_rows.append([f'z{l}',mzz_unmod,max_charge_z])

# Write the data to the CSV file
with open(filename, 'w', newline='') as csvfile:
    writer = csv.writer(csvfile)
    writer.writerows(rows)         

# Generate mod_rows based on specified ranges
for i in range(1, string_length):
    residue = first_line[i-1]
    b_res_num = str(i)
    b_number = static_mods.get(b_res_num) 
    frag_seq_b = first_line[:i]
    max_charge_b_mod = frag_seq_b.count('K') + frag_seq_b.count('H') + frag_seq_b.count('R') +1 -num_charge_neut_nterm
            
    if b_number is not None:
        b_numb_flot = float(b_number)
        if residue in residue_masses:
            mzb_mod = b_numb_flot+ mzb_mod + residue_masses[residue]
    else:
        if residue in residue_masses:
            mzb_mod = mzb_mod + residue_masses[residue]
    mod_rows.append([f'b{i}',mzb_mod, max_charge_b_mod])
    
    
for j in range(1, string_length):
    residue = first_line[j-1]
    c_res_num = str(j)
    c_number = static_mods.get(c_res_num)
    frag_seq_c = first_line[:j]
    max_charge_c_mod = frag_seq_c.count('K') + frag_seq_c.count('H') + frag_seq_c.count('R') +1 -num_charge_neut_nterm
    
    if c_number is not None:
        c_numb_flot = float(c_number)
        if residue in residue_masses:
            mzc_mod = c_numb_flot + mzc_mod + residue_masses[residue]
    else:
        if residue in residue_masses:
            mzc_mod = mzc_mod + residue_masses[residue]
    next_residue = first_line[j]
    if next_residue == "P":
        mod_rows.append([f'c{j}','', max_charge_c_mod])
    else:
        mod_rows.append([f'c{j}',mzc_mod, max_charge_c_mod])

for k in range(1, string_length):
    residue = first_line[string_length-k]
    y_res_num = str(string_length+1-k)
    y_number = static_mods.get(y_res_num)
    backward_seq = ''.join(reversed(first_line))
    frag_seq_y = backward_seq[:k]
    max_charge_y_mod = frag_seq_y.count('K') + frag_seq_y.count('H') + frag_seq_y.count('R') +1 -num_charge_neut
    
    if y_number is not None:
        y_numb_flot = float(y_number)
        if residue in residue_masses:
            mzy_mod = mzy_mod + y_numb_flot + residue_masses[residue]
    else:
        if residue in residue_masses:
            mzy_mod = mzy_mod + residue_masses[residue]
    mod_rows.append([f'y{k}',mzy_mod, max_charge_y_mod])
    
for l in range(1, string_length):
    residue = first_line[string_length-l]
    z_res_num = str(string_length+1-l)
    z_number = static_mods.get(z_res_num)
    frag_seq_z = backward_seq[:l]
    max_charge_z_mod = frag_seq_z.count('K') + frag_seq_z.count('H') + frag_seq_z.count('R') +1 -num_charge_neut
    
    if z_number is not None:
        z_numb_flot = float(z_number)
        if residue in residue_masses:
            mzz_mod = mzz_mod + z_numb_flot + residue_masses[residue]
    else:
        if residue in residue_masses:
            mzz_mod = mzz_mod + residue_masses[residue]
    if residue == 'P':
        mod_rows.append([f'z{l}','', max_charge_z_mod])
    else:
        mod_rows.append([f'z{l}',mzz_mod, max_charge_z_mod])     

# Generate no statics mod_rows based on specified ranges
mzb_mod = proton_mass + float(second_line)
mzc_mod = NH4_mass + float(second_line)
mzy_mod = H3O_mass + float(second_line)
mzz_mod = H2O_minus_NH_mass + float(second_line)

for i in range(1, string_length):
    residue = first_line[i-1]
    b_res_num = str(i)
    b_number = 0 
    frag_seq_b = first_line[:i]
    max_charge_b_mod = frag_seq_b.count('K') + frag_seq_b.count('H') + frag_seq_b.count('R') +1 -num_charge_neut_nterm
            
    if b_number is not None:
        b_numb_flot = float(b_number)
        if residue in residue_masses:
            mzb_mod = b_numb_flot+ mzb_mod + residue_masses[residue]
    else:
        if residue in residue_masses:
            mzb_mod = mzb_mod + residue_masses[residue]
    no_statics_mod_rows.append([f'b{i}',mzb_mod, max_charge_b_mod])
    
    
for j in range(1, string_length):
    residue = first_line[j-1]
    c_res_num = str(j)
    c_number = 0
    frag_seq_c = first_line[:j]
    max_charge_c_mod = frag_seq_c.count('K') + frag_seq_c.count('H') + frag_seq_c.count('R') +1 -num_charge_neut_nterm
    
    if c_number is not None:
        c_numb_flot = float(c_number)
        if residue in residue_masses:
            mzc_mod = c_numb_flot + mzc_mod + residue_masses[residue]
    else:
        if residue in residue_masses:
            mzc_mod = mzc_mod + residue_masses[residue]
    next_residue = first_line[j]
    if next_residue == "P":
        no_statics_mod_rows.append([f'c{j}','', max_charge_c_mod])
    else:
        no_statics_mod_rows.append([f'c{j}',mzc_mod, max_charge_c_mod])

for k in range(1, string_length):
    residue = first_line[string_length-k]
    y_res_num = str(string_length+1-k)
    y_number = 0
    backward_seq = ''.join(reversed(first_line))
    frag_seq_y = backward_seq[:k]
    max_charge_y_mod = frag_seq_y.count('K') + frag_seq_y.count('H') + frag_seq_y.count('R') +1 -num_charge_neut
    
    if y_number is not None:
        y_numb_flot = float(y_number)
        if residue in residue_masses:
            mzy_mod = mzy_mod + y_numb_flot + residue_masses[residue]
    else:
        if residue in residue_masses:
            mzy_mod = mzy_mod + residue_masses[residue]
    no_statics_mod_rows.append([f'y{k}',mzy_mod, max_charge_y_mod])
    
for l in range(1, string_length):
    residue = first_line[string_length-l]
    z_res_num = str(string_length+1-l)
    z_number = 0
    frag_seq_z = backward_seq[:l]
    max_charge_z_mod = frag_seq_z.count('K') + frag_seq_z.count('H') + frag_seq_z.count('R') +1 -num_charge_neut
    
    if z_number is not None:
        z_numb_flot = float(z_number)
        if residue in residue_masses:
            mzz_mod = mzz_mod + z_numb_flot + residue_masses[residue]
    else:
        if residue in residue_masses:
            mzz_mod = mzz_mod + residue_masses[residue]
    if residue == 'P':
        no_statics_mod_rows.append([f'z{l}','', max_charge_z_mod])
    else:
        no_statics_mod_rows.append([f'z{l}',mzz_mod, max_charge_z_mod])   

def not_within_range(value, num_list, range_value):
    for num in num_list:
        if abs(num - value) <= range_value:
            return False
    return True
    
# Write the data to the CSV file
with open(filename1, 'w', newline='') as csvfile:
    writer = csv.writer(csvfile)
    writer.writerows(mod_rows)
    
with open(filename2, 'w', newline='') as csvfile:
    writer = csv.writer(csvfile)
    writer.writerows(no_statics_rows)
    
with open(filename3, 'w', newline='') as csvfile:
    writer = csv.writer(csvfile)
    writer.writerows(no_statics_mod_rows)


def compare_unmod_csv_files(unmod_potential_file, unmod_data_file, matched, matched_for_qpi_unmod, matched_for_qpi_mod, mod_potential_file, mod_data_file):
    matched = []
    matched_for_qpi_unmod = []
    matched_for_qpi_mod = []
    
  # Read unmod_potential file into a dictionary
    with open(unmod_potential_file, 'r') as unmod_potential_csv:
        unmod_potential_reader = csv.reader(unmod_potential_csv)
        unmod_potential_dict = { q[0]:q[1] for q in unmod_potential_reader }
    with open(unmod_potential_file, 'r') as unmod_potential_csv:
        unmod_potential_reader = csv.reader(unmod_potential_csv)
        unmod_potential_dict2 = { r[0]:r[2] for r in unmod_potential_reader }
    
    # Read unmod_data file and perform comparison
    with open(unmod_data_file, 'r') as unmod_data_csv:
        unmod_data_reader = csv.reader(unmod_data_csv)
        next(unmod_data_reader)  # Skip header
        for row in unmod_data_reader:
            if row[1] in ['a','b','c']:
                key = row[1] + row[2]
            elif row[1] in ['y','z']:
                seq = string_length - int(row[2])
                key = row[1] + str(seq)
               
            if key in unmod_potential_dict:
                    if (unmod_potential_dict[key]) != '':
                        unmod_potential_value = float(unmod_potential_dict[key])
                    if (unmod_potential_dict2[key]) != '':
                        unmod_potential_charge = int(unmod_potential_dict2[key])
                    measured_charge = int(row[0])
                    
                    if unmod_potential_charge >= measured_charge:
                    
                        
                    
                        if row[1][0] == 'b':
                            unmod_calculated_value = float(row[0]) * float(row[3]) - (float(row[0]) - 1) * 1.00728
                            if abs(unmod_calculated_value - unmod_potential_value) <= 0.5:
                                matched.append([unmod_potential_value - 1.00728, float(row[5])])
                                matched_for_qpi_unmod.append(key)
                            else:
                                with open('log.txt', 'a') as log_file:
                                    log_file.write(key + ' ' + row[0] + '+: Incorrect identification in EXDViewer, measured monoisotopic m/z was ' + str(unmod_calculated_value) + ' and theoretical monoisotopic m/z is ' + str(unmod_potential_value) + '. Removed from ' + unmod_data_file + '\n')
                        elif row[1][0] == 'c':
                            unmod_calculated_value = float(row[0]) * float(row[3]) - (float(row[0]) - 1) * 1.00728
                            if abs(unmod_calculated_value - unmod_potential_value) <= 0.5:
                                 matched.append([unmod_potential_value - 1.00728, float(row[5])])
                                 matched_for_qpi_unmod.append(key)
                            elif unmod_potential_value - unmod_calculated_value <= 2.5:
                                 if unmod_potential_value >= unmod_calculated_value:
                                     matched.append([unmod_potential_value - 1.00728, float(row[5])])
                                     matched_for_qpi_unmod.append(key)
                            else:
                                with open('log.txt', 'a') as log_file:
                                    log_file.write(key + ' ' + row[0] + '+: Incorrect identification in EXDViewer, measured monoisotopic m/z was ' + str(unmod_calculated_value) + ' and theoretical monoisotopic m/z is ' + str(unmod_potential_value) + '. Removed from ' + unmod_data_file + '\n')
                                    
                        elif row[1][0] == 'y':
                           
                             unmod_calculated_value = float(row[0]) * float(row[3]) - (float(row[0]) - 1) * 1.00728
                             if abs(unmod_calculated_value - unmod_potential_value) <= 0.5:
                                 matched.append([unmod_potential_value - 1.00728, float(row[5])])
                                 matched_for_qpi_unmod.append(key)
                             else:
                                 with open('log.txt', 'a') as log_file:
                                     log_file.write(key + ' ' + row[0] + '+: Incorrect identification in EXDViewer, measured monoisotopic m/z was ' + str(unmod_calculated_value) + ' and theoretical monoisotopic m/z is ' + str(unmod_potential_value) + '. Removed from ' + unmod_data_file + '\n')
                                 
                        elif row[1][0] == 'z':
                             unmod_calculated_value = float(row[0]) * float(row[3]) - (float(row[0]) - 1) * 1.00728
                             if abs(unmod_calculated_value - unmod_potential_value) <= 0.5:
                                 matched.append([unmod_potential_value - 1.00728, float(row[5])])
                                 matched_for_qpi_unmod.append(key)
                             elif unmod_calculated_value - unmod_potential_value <= 2.5:
                                 if unmod_calculated_value >= unmod_potential_value:
                                     matched.append([unmod_potential_value - 1.00728, float(row[5])])
                                     matched_for_qpi_unmod.append(key)
                             else:
                                 with open('log.txt', 'a') as log_file:
                                    log_file.write(key + ' ' + row[0] + '+: Incorrect identification in EXDViewer, measured monoisotopic m/z was ' + str(unmod_calculated_value) + ' and theoretical monoisotopic m/z is ' + str(unmod_potential_value) + '. Removed from ' + unmod_data_file + '\n')
                    else:
                        with open('log.txt', 'a') as log_file:
                            log_file.write(key + ' ' + row[0] + '+: Charge is higher than maximum charge ' + str(unmod_potential_charge) + '+, removed from ' + unmod_data_file + '\n')
                            
    unmod_masses = [sublist[0] for sublist in matched]
  
    
    # Read mod potential file into the dictionary
    with open(mod_potential_file, 'r') as mod_potential_csv:
        mod_potential_reader = csv.reader(mod_potential_csv)
        mod_potential_dict = { q[0]:q[1] for q in mod_potential_reader }
    with open(mod_potential_file, 'r') as mod_potential_csv:
        mod_potential_reader = csv.reader(mod_potential_csv)
        mod_potential_dict2 = { r[0]:r[2] for r in mod_potential_reader }
    
    # Read mod_data file and perform comparison
    with open(mod_data_file, 'r') as mod_data_csv:
        mod_data_reader = csv.reader(mod_data_csv)
        next(mod_data_reader)  # Skip header
        for row in mod_data_reader:
            if row[1] in ['a','b','c']:
                key = row[1] + row[2]
            elif row[1] in ['y','z']:
                seq = string_length - int(row[2])
                key = row[1] + str(seq)
                
            if key in mod_potential_dict:
                if (mod_potential_dict[key]) != '':
                    mod_potential_value = float(mod_potential_dict[key])
                mod_calculated_value = float(row[0]) * float(row[3]) - (float(row[0]) - 1) * 1.00728
                mod_calculated_mass = mod_calculated_value - 1.00728
                if (mod_potential_dict2[key]) != '':
                   mod_potential_charge = int(mod_potential_dict2[key])
                measured_charge = int(row[0])
                
                if mod_potential_charge >= measured_charge:           
                    if not_within_range(mod_calculated_mass, unmod_masses, 2.5):
                    
                            if row[1][0] == 'b': 
                                            if abs(mod_calculated_value - mod_potential_value) <= 0.5:
                                                matched.append([mod_potential_value - 1.00728, float(row[5])])
                                                matched_for_qpi_mod.append(key)
                                            else:
                                                with open('log.txt', 'a') as log_file:
                                                    log_file.write(key + ' ' + row[0] + '+: Incorrect identification in EXDViewer, measured monoisotopic m/z was ' + str(mod_calculated_value) + ' and theoretical monoisotopic m/z is ' + str(mod_potential_value) + '. Removed from ' + mod_data_file + '\n')
                            elif row[1][0] == 'c':
                                            if abs(mod_calculated_value - mod_potential_value) <= 0.5:
                                                matched.append([mod_potential_value - 1.00728, float(row[5])])
                                                matched_for_qpi_mod.append(key)
                                            elif mod_potential_value - mod_calculated_value <= 2.5:
                                                if mod_potential_value >= mod_calculated_value:
                                                    matched.append([mod_potential_value - 1.00728, float(row[5])])
                                                    matched_for_qpi_mod.append(key)
                                            else:
                                                with open('log.txt', 'a') as log_file:
                                                   log_file.write(key + ' ' + row[0] + '+: Incorrect identification in EXDViewer, measured monoisotopic m/z was ' + str(mod_calculated_value) + ' and theoretical monoisotopic m/z is ' + str(mod_potential_value) + '. Removed from ' + mod_data_file + '\n')
                            elif row[1][0] == 'y':
                                            if abs(mod_calculated_value - mod_potential_value) <= 0.5:
                                                matched.append([mod_potential_value - 1.00728, float(row[5])])
                                                matched_for_qpi_mod.append(key)
                                            else:
                                                with open('log.txt', 'a') as log_file:
                                                   log_file.write(key + ' ' + row[0] + '+: Incorrect identification in EXDViewer, measured monoisotopic m/z was ' + str(mod_calculated_value) + ' and theoretical monoisotopic m/z is ' + str(mod_potential_value) + '. Removed from ' + mod_data_file + '\n')
                            elif row[1][0] == 'z':
                                            if abs(mod_calculated_value - mod_potential_value) <= 0.5:
                                                matched.append([mod_potential_value - 1.00728, float(row[5])])
                                                matched_for_qpi_mod.append(key)
                                            elif mod_calculated_value - mod_potential_value <= 2.5:
                                                if mod_calculated_value >= mod_potential_value:
                                                    matched.append([mod_potential_value - 1.00728, float(row[5])])
                                                    matched_for_qpi_mod.append(key)
                                            else:
                                                with open('log.txt', 'a') as log_file:
                                                   log_file.write(key + ' ' + row[0] + '+: Incorrect identification in EXDViewer, measured monoisotopic m/z was ' + str(mod_calculated_value) + ' and theoretical monoisotopic m/z is ' + str(mod_potential_value) + '. Removed from ' + mod_data_file + '\n')

                else:
                    with open('log.txt', 'a') as log_file:
                        log_file.write(key + ' ' + row[0] + '+: Charge is higher than maximum charge ' + str(mod_potential_charge) + '+, removed from ' + mod_data_file + '\n')

    return matched, matched_for_qpi_unmod, matched_for_qpi_mod

def remove_duplicates(nonmod_file, mod_file, output_file):
    with open(nonmod_file, 'r') as nonmod_csv:
        nonmod_reader = csv.reader(nonmod_csv)
        nonmod_lines = set(tuple(row[:3]) for idx, row in enumerate(nonmod_reader) if idx != 0)

    mod_lines = []    
    with open(mod_file, 'r') as mod_csv:
        mod_reader = csv.reader(mod_csv)
        header = next(mod_reader)  # Store the header row
        for idx, row in enumerate(mod_reader):
            if tuple(row[:3]) in nonmod_lines:
                # Write duplicate line to log file
                with open('log.txt', 'a') as log_file:
                    if row[1] in ['a','b','c']:
                        log_file.write(row[1] + row[2] + ' ' + row[0] + '+: Duplicate found in nonmod, removed from ' + mod_file + '\n')
                    elif row[1] in ['y','z']:
                        corrected_ind = string_length - int(row[2])
                        log_file.write(row[1] + str(corrected_ind) + ' ' + row[0] + '+: Duplicate found in nonmod, removed from ' + mod_file + '\n')
            else:
                mod_lines.append(row)


    with open(output_file, 'w', newline='') as output_csv:
        writer = csv.writer(output_csv)
        writer.writerows(mod_lines)


def delete_duplicate_lines(filename):
  """Deletes duplicate lines from a text file.

  Args:
    filename: The name of the text file.
  """

  # Create a set to store the unique lines.
  unique_lines = set()

  # Open the file for reading.
  with open(filename, "r") as f:
    # Read each line in the file.
    for line in f:
      # If the line is not in the set, add it.
      if line not in unique_lines:
        unique_lines.add(line)

  # Open the file for writing.
  with open(filename, "w") as f:
    # Write each line in the set to the file.
    for line in unique_lines:
      f.write(line)

# Get the values for QPI; must remove static mod masses just to make the software work
def get_qpi_masses(unmod_list, mod_list, unmod, mod, unmod_int, mod_int):
   
   unmod_tuple = tuple(unmod_list)
   mod_tuple = tuple(mod_list)
   mass_int_pairs = []
   with open(unmod, "r") as f:
        mass_reader = csv.reader(f)
        fragid = []
        nostaticmass = []
        for row in mass_reader:
           fragid.append(row[0])
           nostaticmass.append(row[1])
   mass_dict = dict(zip(fragid,nostaticmass))
   
   with open(unmod_int, "r") as g:
        letters = []
        numbers = []
        intense = []
        intensity_reader = csv.reader(g)
        next(intensity_reader)
        for row in intensity_reader:
            for row in intensity_reader:
              letters.append(row[1])
              if row[1] in ['a','b','c']:
                  numbers.append(row[2])
              elif row[1] in ['y','z']:
                  numbers.append(str(string_length - int(row[2])))
              intense.append(row[5])
            
    
   frag_name = [x + y for x, y in zip(letters, numbers)]
   intense_dict = dict(zip(frag_name, intense))
   intense_dict = merge_duplicate_keys(intense_dict)


   for item in unmod_tuple:
       if item in mass_dict:
           if item in intense_dict:
               m = str(float(mass_dict[item]) - 1.00728)
               i = intense_dict[item]
               mi = [m,i]
               mass_int_pairs.append(mi)


#make this like the second to return a dictionary of masses with keys as fragment name
   with open(mod, "r") as f:
        mass_reader = csv.reader(f)
        fragid = []
        nostaticmass = []
        for row in mass_reader:
            fragid.append(row[0])
            nostaticmass.append(row[1])
   mod_mass_dict = dict(zip(fragid,nostaticmass))
        
   with open(mod_int, "r") as g:
       letters = []
       numbers = []
       intense = []
       intensity_reader = csv.reader(g)
     
       for row in intensity_reader:
         letters.append(row[1])
         if row[1] in ['a','b','c']:
             numbers.append(row[2])
         elif row[1] in ['y','z']:
             numbers.append(str(string_length - int(row[2])))
         intense.append(row[5])

   mod_frag_name = [x + y for x, y in zip(letters, numbers)]
   mod_intense_dict = dict(zip(mod_frag_name, intense))
   mod_intense_dict = merge_duplicate_keys(mod_intense_dict)


   for item in mod_tuple:
        if item in mod_mass_dict:
            if item in mod_intense_dict:
                m = str(float(mod_mass_dict[item]) - 1.00728)
                i = mod_intense_dict[item]
                mi = [m,i]
                mass_int_pairs.append(mi)


   return mass_int_pairs      
        
   


# Comparing data to theoretical values and writing csv files for QPI and removing IDs from mod that appear in nonmod (i.e., false positives)
unmod_potential_file = 'potential_unmods.csv'
unmod_data_file = 'unmod_rep1.csv'
mod_potential_file = 'potential_mods.csv'
mod_data_file = 'mod_rep1.csv'
matched = 'rep1_matched'
matched_for_qpi_unmod = 'rep1_qpi_matched_unmod'
matched_for_qpi_mod = 'rep1_qpi_matched_mod'
no_statics_unmod_file = 'no_statics_unmods.csv'
no_statics_mod_file = 'no_statics_mods.csv'
sum_intensity_isotopes(unmod_data_file)
sum_intensity_isotopes(mod_data_file)
remove_duplicates('nonmod_rep1.csv', 'mod_rep1.csv', 'no_fp_mod_rep1.csv')
mod_data_file = 'no_fp_mod_rep1.csv'
rep1_matched = compare_unmod_csv_files(unmod_potential_file, unmod_data_file, matched, matched_for_qpi_unmod, matched_for_qpi_mod, mod_potential_file, mod_data_file)
summed_duplicates_rep1 = process_list(rep1_matched[0])
print('The following modified ions were matched in rep1: ')
for item in rep1_matched[2]:
     print(item)
write_deconv_qpi_file(summed_duplicates_rep1, 'rep1_masslist')
rep1_for_qpi = get_qpi_masses(rep1_matched[1], rep1_matched[2], no_statics_unmod_file, no_statics_mod_file, unmod_data_file, mod_data_file)
write_deconv_qpi_file(rep1_for_qpi, 'rep1')

# Comparing data to theoretical values and writing csv files for QPI and removing IDs from mod that appear in nonmod (i.e., false positives)
unmod_potential_file = 'potential_unmods.csv'
unmod_data_file = 'unmod_rep2.csv'
mod_potential_file = 'potential_mods.csv'
mod_data_file = 'mod_rep2.csv'
matched = 'rep2_matched'
matched_for_qpi_unmod = 'rep2_qpi_matched_unmod'
matched_for_qpi_mod = 'rep2_qpi_matched_mod'
no_statics_unmod_file = 'no_statics_unmods.csv'
no_statics_mod_file = 'no_statics_mods.csv'
sum_intensity_isotopes(unmod_data_file)
sum_intensity_isotopes(mod_data_file)
remove_duplicates('nonmod_rep2.csv', 'mod_rep2.csv', 'no_fp_mod_rep2.csv')
mod_data_file = 'no_fp_mod_rep2.csv'
rep2_matched = compare_unmod_csv_files(unmod_potential_file, unmod_data_file, matched, matched_for_qpi_unmod, matched_for_qpi_mod, mod_potential_file, mod_data_file)
summed_duplicates_rep2 = process_list(rep2_matched[0])
print('The following modified ions were matched in rep2: ')
for item in rep2_matched[2]:
     print(item)
write_deconv_qpi_file(summed_duplicates_rep2, 'rep2_masslist')
rep2_for_qpi = get_qpi_masses(rep2_matched[1], rep2_matched[2], no_statics_unmod_file, no_statics_mod_file, unmod_data_file, mod_data_file)
write_deconv_qpi_file(rep2_for_qpi, 'rep2')

# Comparing data to theoretical values and writing csv files for QPI and removing IDs from mod that appear in nonmod (i.e., false positives)
unmod_potential_file = 'potential_unmods.csv'
unmod_data_file = 'unmod_rep3.csv'
mod_potential_file = 'potential_mods.csv'
mod_data_file = 'mod_rep3.csv'
matched = 'rep3_matched'
matched_for_qpi_unmod = 'rep3_qpi_matched_unmod'
matched_for_qpi_mod = 'rep3_qpi_matched_mod'
no_statics_unmod_file = 'no_statics_unmods.csv'
no_statics_mod_file = 'no_statics_mods.csv'
sum_intensity_isotopes(unmod_data_file)
sum_intensity_isotopes(mod_data_file)
remove_duplicates('nonmod_rep3.csv', 'mod_rep3.csv', 'no_fp_mod_rep3.csv')
mod_data_file = 'no_fp_mod_rep3.csv'
rep3_matched = compare_unmod_csv_files(unmod_potential_file, unmod_data_file, matched, matched_for_qpi_unmod, matched_for_qpi_mod, mod_potential_file, mod_data_file)
summed_duplicates_rep3 = process_list(rep3_matched[0])
print('The following modified ions were matched in rep3: ')
for item in rep3_matched[2]:
     print(item)
write_deconv_qpi_file(summed_duplicates_rep3, 'rep3_masslist')
rep3_for_qpi = get_qpi_masses(rep3_matched[1], rep3_matched[2], no_statics_unmod_file, no_statics_mod_file, unmod_data_file, mod_data_file)
write_deconv_qpi_file(rep3_for_qpi, 'rep3')


def compare_csv_files(csv_files, output_file):
    # Create a set to store values
    values_set = set()

    # Iterate through each CSV file
    for csv_file in csv_files:
        with open(csv_file, 'r') as file:
            csv_reader = csv.reader(file)
            # Skip header if present
            next(csv_reader, None)
            # Iterate through rows
            for row in csv_reader:
                # Check if the first item of the row is in values_set
                if row[0] in values_set:
                    # If found in at least two files, write to output file
                    with open(output_file, 'a') as out_file:
                        out_file.write(row[0] + '\n')
                else:
                    # Otherwise, add it to the set
                    values_set.add(row[0])

if __name__ == "__main__":
    # List of CSV files to compare
    csv_files = ['rep1_masslist.csv', 'rep2_masslist.csv', 'rep3_masslist.csv']
    # Output file to write matching values
    output_file = 'matched_masses.txt'

    # Clear content of output file if it already exists
    with open(output_file, 'w'):
        pass

    # Compare CSV files
    compare_csv_files(csv_files, output_file)

delete_duplicate_lines("log.txt")
